# Hypothesis-Driven Identification of Neural Algorithms With Dynamical Structure-Preserving Manifolds

**DOI:** 10.64898/2026.10.09.755459

**Authors:** Daniel Calbick, Jason Z. Kim, Hansem Sohn, Ilker Yildirim

## Abstract

Neuroscience aims to uncover the algorithms by which the brain builds and manipulates rich internal representations of objects, agents, and places. Existing modeling frameworks have had limited success in identifying these algorithms. Here, we present Dynamical Structure-Preserving Manifolds (dSPMs), a multilevel framework that programs, without training, recurrent neural networks to enable hypothesis-driven identification of the brain’s algorithms that generate symbolic, structure-preserving representations of the world. We apply dSPM to identify an algorithm for physical scene prediction in the dorsomedial frontal cortex (DMFC) of macaques intercepting a ball on a Pong-like virtual gameboard. dSPM better explained DMFC than alternatives, made predictions confirmed in neural data, and revealed an algorithm of DMFC that we term ‘manifold-based simulation’: perceived initial conditions rapidly collapse neural activity onto a low-dimensional manifold whose geometry encodes a physics-based representation of the scene, supporting far-out predictions without traditional next-step simulation. dSPM supports effective exploration of the computational foundations of biological intelligence.

## Introduction

The brain generates rich internal representations of the world around us, enabling predictions about how objects will move, agents will act, and places will connect. A central challenge of neuroscience is to uncover the algorithms — i.e., the precise, if possible interpretable, computational transformations — by which the brain builds and manipulates these complex mental representations.

Two computational modeling approaches to identifying the brain’s algorithms dominate current work, each focusing on different perspectives on cognition and offering distinct advantages and limitations. On one hand, probabilistic models of cognition specify interpretable hypotheses about mental representations [1], but typically lack neural grounding. These models suggest that the brain involves approximate structure-preserving representations of the external data-generating processes [2] — e.g., the physics that govern how objects move and react to external forces — that in turn support predictions of what will happen next [3–5]. Functionally, structure-preserving representations are appealing: Having access to the worldly causes underlying a scene is behaviorally efficacious [6], supporting efficient learning, flexible generalization [7–9], and compositionality [10]. However, these hypotheses are typically formulated as “computational-level” accounts of the system [11, 12]: i.e., in practice, they are implemented using standard computer software and high-level programming languages, without identifying an actual algorithm that could realize them in biological neural circuits. On the other hand, deep neural network (DNN) models fit some aspects of neural data but obscure the algorithms they learn. DNNs facilitate explorations of task-optimized statistical representations as accounts of mental representations [13–15], which can involve “shortcuts” and “simple tricks” that are typically, but not always, performant [15, 16]. Crucially, DNNs offer scientific hypotheses with respect to their architectures, training data, and training objectives [17], and not directly with respect to the possible algorithms themselves. Instead, these “implicit algorithms” need to be “extracted” after the fact using various interpretation tools, regardless of whether these DNNs are large or small (e.g., [18, 19]).

This landscape of modeling approaches highlights a critical gap: intuitive cognitive theories that are difficult to map to neurobiology, and fits to neural data that do not allow algorithmic-level control over hypothesis testing. A new approach is needed to precisely and interpretably identify algorithms of complex mental representations in the brain.

Here, we introduce Dynamical Structure-Preserving Manifolds (dSPM) for hypothesis-driven identification of neural algorithms and present a case study of physical scene prediction in macaque frontal circuitry to establish its efficacy. At its core, dSPM unifies two influential but, to this day, disparate views of the brain that cut across the fields of cognitive science and neuroscience: that the brain can be viewed as a symbolic representation system building and manipulating approximate structure-preserving maps of the external data-generating processes [2, 20] (Fig. 1A) and that the brain can be viewed as the evolution of coupled dynamical systems in population state space [21, 22] (Fig. 1B). Even though dynamical systems offer a common framework for neuroscience, they are most often used as an interpretive tool to analyze and visualize the otherwise high-dimensional activity (e.g., [23, 24]). The dSPM framework’s “top-down” construction of computation through dynamics, by specifying dynamical algorithms that realize complex symbolic representations, contrasts with this traditional approach.

**Figure 1.**
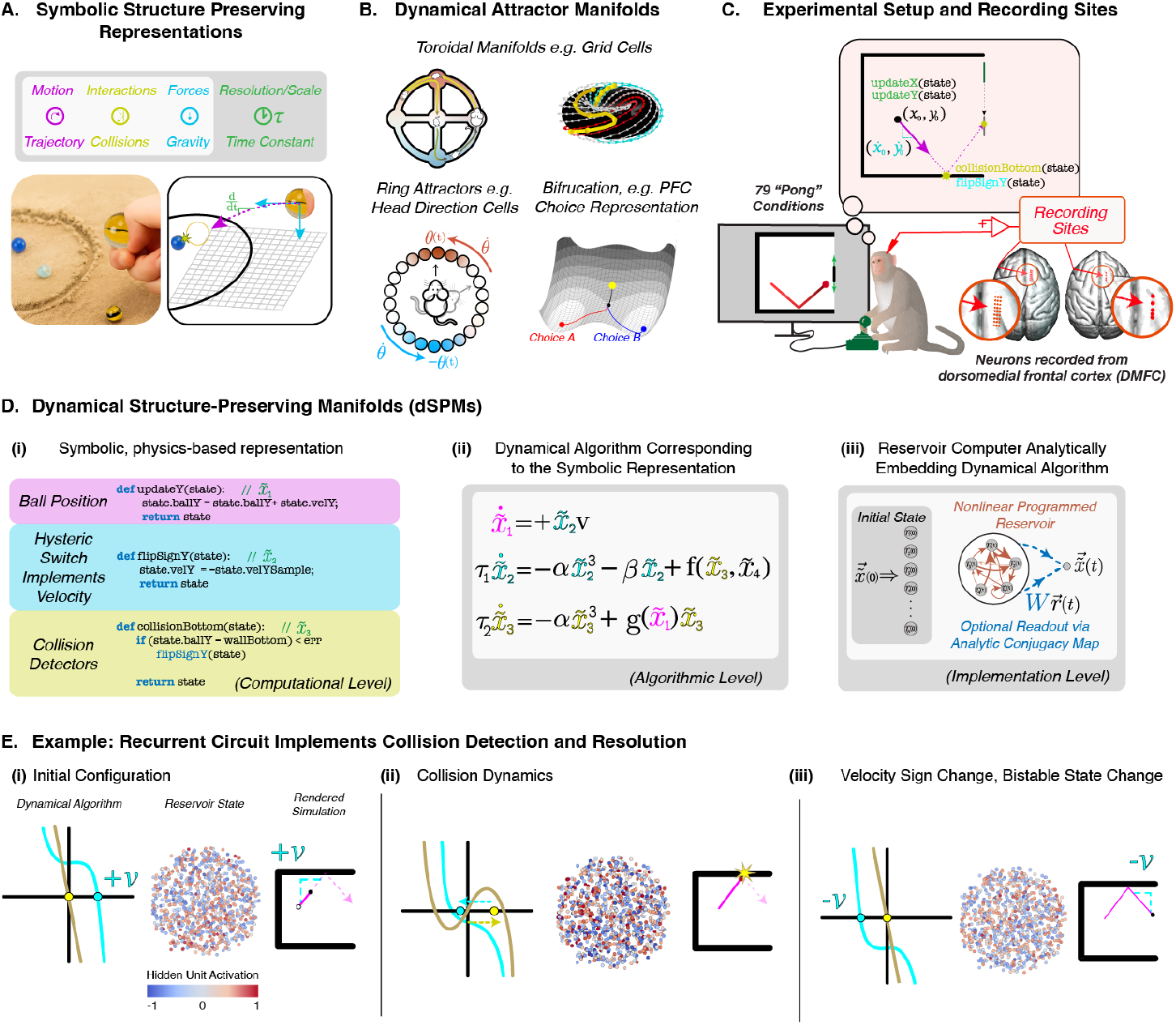
Overview of the Dynamical Structure-Preserving Manifolds framework. (A) The brain can be viewed as a representation system with approximate structure-preserving mappings of the external data-generating processes, including objects and the physics that govern their movement. (B) Example dynamical systems in neuroscience implementing complex cognitive representations, including a ring attractor for heading direction, a torus for position in 2D space, and bifurcation for decision-making. (C) Experimental design showing neural recordings from DMFC in two macaques performing a ball interception task in a Pong-like gameboard. The “thought bubble” indicates the hypothesis that dSPM enables us to test, ‘manifold-based simulation’: DMFC circuitry builds a physics-based representation of the gameboard, including the entities and relations of the ball, walls, and the paddle, embedded in a low-dimensional manifold. (D-i) At the computational level, the dSPM framework posits that predictions are derived from a physics-based representation of the gameboard, e.g., a posterior predictive distribution over future trajectory conditioned on observed trajectory. Typical cognitive models implement this objective using standard computer software and high-level programming languages, without any neural grounding. (D-ii) The dSPM’s dynamical algorithm for the Pong setting, i.e., a set of coupled differential equations translating the program-like representation in panel (D-i). Further explained in panel (E). (D-iii) The dSPM’s reservoir computer analytically embeds the dynamical algorithm in its connectivity matrix, without training, a training objective, or a training dataset. This reservoir computer is a neurally falsifiable instantiation of the physics-based representation hypothesis. (E) Collision detection and resolution in dSPM through dynamics and recurrent circuits that undergo phase transitions when the ball approaches a wall. In each subpanel, the leftmost plots show the phase portrait of the two coupled dynamical variables (out of 12) (cyan: velocity sign, 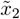; yellow: collision detection, 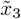); the middle scatterplots show the hidden units of the reservoir computer; the rightmost panels visualize the gameboard. (E-i) Unfolding the initial configuration by integrating a positive velocity value; (E-ii) Collision detection and resolution via cubic bifurcation (yellow line), which non-linearly bifurcates from one stable attractor in (E-i) to two stable attractors (outer zero-crossings) and one repeller (middle zero-crossing) due to proximity to the wall. This transition pulls the velocity variable, causing a sign flip (blue line). (E-iii) Unfolding the rest of the simulation, similar to (E-i), but by integrating this negative velocity value −*v*.

The dSPM framework achieves this in two stages. First, a computational hypothesis, typically expressed symbolically in cognitive modeling (e.g., future trajectory prediction implemented in a high-level programming language), is formulated as a dynamical algorithm. The dynamical algorithm creates a computational graph, including discrete operations such as if-else branching, realizing the program-like representation through dynamically changing, coupled attractors and repellors. Second, dSPM embeds this dynamical algorithm, typically of low dimensionality, in a much higher-dimensional reservoir computer, a biologically plausible architecture for recurrent, distributed computation [25], by analytically setting the reservoir’s connectivity (i.e., the adjacency matrix) without any training, training data, or training objectives. The result is a population of interconnected units whose analytically set connectivity directs the flow of information to align with the dynamical algorithm of the hypothesized structure-preserving representation.

Accordingly, dSPM provides a multilevel computational modeling tool, filling the modeling gap mentioned earlier: Unlike task-optimized DNNs, dSPM’s reservoir computer is a white-box at the algorithm level (i.e., enabling direct hypothesis-testing of specific algorithms), and unlike probabilistic models of cognition, dSPM is falsifiable at the neural implementation level, enabling hypothesis-driven exploration of the algorithms of the brain’s complex cognitive representations.

To assess the efficacy of dSPM, we apply it to identify an algorithm for predicting physical scenes in macaque frontal circuitry. Rajalingham et al. [26] performed high-throughput single-cell recordings in the dorsomedial frontal cortex (DMFC) of macaques performing a virtual ball interception task in a Pong-like gameboard. The researchers occluded the ball’s trajectory partway through the trial, finding evidence of prediction in the DMFC activity. How does the DMFC population represent the gameboard, and how does this representation support the prediction of future trajectory? The dSPM framework enables the novel capability of testing the hypothesis of a physics-based representation of the gameboard — including positions of the ball and the walls, velocity, and collision state, as well as interaction rules (as in the cognitive modeling of ref. [3]) — directly in distributed, dynamic neural activity.

The dSPM framework instantiated to this domain generates a cascade of falsifiable predictions about DMFC activity. First, the perceived initial conditions of a scene should rapidly, i.e., within trial onset, collapse high-dimensional neural activity onto a compact, low-dimensional manifold — and the geometry of that manifold should reflect not merely the momentary sensory content, but something about the causal structure of the scene. Second, because the collapsed manifold encodes the physics of the gameboard, any single population state on the manifold should be sufficient to linearly decode the entire future trajectory of the ball without the traditional, step-by-step simulation — which we term ‘manifold-based simulation’. Third, the dSPM’s cross-condition representational similarity should substantially and uniquely explain DMFC’s cross-condition similarity structure, beyond what alternative models can account for. We confirm each of these three predictions of dSPM in the DMFC data. In both quantitative and qualitative ways, dSPM outperforms several alternative models, including the statistical representations in a series of task-optimized recurrent neural networks and a task-performant heuristic specialized to this ball-interception task. Accordingly, dSPM reveals an algorithm of DMFC in which a latent, low-dimensional manifold encodes a structure-preserving physics-based representation of physical scenes, enabling future predictions without step-by-step simulation or task-performance heuristics. dSPM offers a general approach to hypothesis-driven, multilevel identification of neural algorithms undergirding the rich representations that make up our mental life.

## Dynamical Structure-Preserving Manifolds

Cognitive theories formalize physical prediction as building and running forward an approximate, structure-preserving representation Ψ of the worldly causes governing how objects move and react to forces (position and velocity of objects, their physical properties, and approximate Newtonian mechanics) and computing a posterior over future trajectories parametrized by Ψ [3, 6] (Fig. 1C “thought bubble”). Existing implementations of Ψ rely on off-the-shelf computer graphics software and high-level programming languages (e.g., [3–5, 27]), and therefore cannot inform how biological neural systems could realize such representations. dSPM closes this gap in two stages: expressing Ψ as a dynamical algorithm, and realizing that algorithm in a neurally falsifiable model class.

The first stage builds on a clue from recent neuroscience: the geometry of low-dimensional latent manifolds underlying neural population activity corresponds to approximate structure-preserving maps of the world [22] (Fig. 1B): e.g., ring attractors for heading direction, toroidal geometry for position in 2D space [28–31]. We propose that approximate structure-preserving representations of physical scenes are likewise encoded in dynamical algorithms latent in high-dimensional neural activity. A dynamical algorithm composes a set of variables 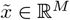 and their flow through time to induce an *M*-dimensional flow field with time-varying attractors (sinks) and repellors (sources) (Supplementary Fig. 5A). Unlike the common fixed-point attractors [32, 33], this coupling defines a ‘dynamical manifold’ whose attractors and repellers not only change over time, but can be computationally harnessed to build sophisticated computational graphs (cf. [34]). This substrate is sufficiently expressive to realize the computational graph motifs of high-level programming languages, such as Boolean logic, if-else branching, and loops [35]; data structures and algorithms, such as sorting, DFT, and memory storage and access [36, 37]; and numerical computation such as optimization, inverse problems, and simulation [35, 38]. Dynamical algorithms are powerful not because they re-express classical algorithms in a different way, but rather, these algorithms, in a fundamental way, bring us to the equivalence class of biologically plausible computation — flow fields induced by populations of nonlinear leaky-integrator neurons.

This takes us to the second stage, which embeds the low-dimensional, symbolic dynamical algorithm into the high-dimensional state space of a neurally mappable model class, a tanh-activated reservoir computer [25,39], by *programming* rather than training. In other words, a high-dimensional, non-linear reservoir with dim(*r*) = *N* hosts a low-dimensional, symbolic algorithm with 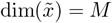 through a topological semi-conjugacy map [40, 41] that translates one non-linear flow to another. To do so, we begin from an open-loop reservoir of leaky integrators,

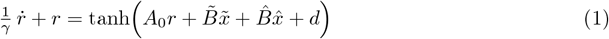

whose inputs are split into the external drives 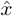 and the open-loop (i.e., to-be-programmed) recurrent variables 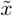 whose time derivatives parametrize the target flow field (i.e., the dynamical algorithm). Following a recent advance in the physics of dynamical systems [35], we “decompile” this reservoir, expanding a closed-form solution of its hidden state *r*(*t*) into a polynomial basis *ϕ* consisting of the open-loop variables and their time derivatives, *r*(*t*) ≈ *R ϕ*, and solve for the readout *W* that creates a map between the decompiled reservoir state and the target dynamical algorithm written in the same basis:

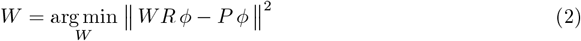

where *R* and *P* are the coefficient matrices of the decompiled reservoir state and the target dynamical algorithm in the shared basis *ϕ*, respectively. Because *R ϕ* stands for the reservoir state itself, the solved readout reproduces the target variables from that state, 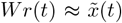. Closing the loop through this readout 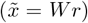 yields an autonomous reservoir whose connectivity matrix *A*_*p*_ is programmed:

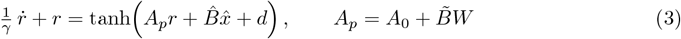

We emphasize what this construction does and does not entail. First, the reservoir’s activity is an analytical consequence of the hypothesized Ψ, thereby providing a computational theory specified across multiple levels of analysis. Second, although the programmed dynamical algorithm is symbolic, what is actually run is nothing symbolic at all: a fully nonlinear dynamical system of leaky integrators with tanh saturation whose recurrent connections are strong far from the weakly coupled regime of conventional reservoir computing [42] (see Methods and Supplementary Section 3 for a spectral analysis). Consequently, all subsequent comparisons between dSPM and DMFC should be read in this light: two high-dimensional populations of nonlinear leaky integrators, each evolving under its own recurrent dynamics.

In Supplementary Section 2 and Supplementary Figs. 1 and 2, we illustrate dSPM’s expressivity with a programmed Lotka–Volterra system [43, 44], whose conserved Hamiltonian provides an exact benchmark: built with our MATLAB interface and without training, the programmed reservoir sustains genuinely nonlinear, conservative dynamics.

### Applying dSPM to a ball interception task

To test dSPM for identifying a neural algorithm of physical scene prediction, we leveraged a virtual ball-interception task, developed by Rajalingham et al. [45, 46]. In this task, rhesus macaques (*Macaca mulatta*) used a joystick to control a paddle to intercept a moving ball on a computer screen (Fig. 1C), much like the classical video game of Pong. During the latter portion of its trajectory, the ball becomes occluded, requiring some prediction strategy for successful interception. The task design systematically varies the initial position and velocity of the ball across 79 unique conditions, yielding a range of trajectories with different durations (range: 1450-3750 ms). Each condition ends when the ball reaches the rightmost side of the board, whether intercepted or not. Large-scale electrophysiological recordings from the dorsomedial frontal cortex (DMFC) of two macaques performing this task yielded activity from 1,889 neurons. The DMFC population activity contained information about the ball position regardless of whether it was occluded, making it a prime candidate for implementing physical scene prediction [46].

To test the hypothesis depicted in Fig. 1C, we start by symbolically expressing (i.e., using a high-level programming language) a physics-based representation of the gameboard Ψ, including the walls, paddle, ball, the ball’s position and how it changes over time (linear dynamics), if-else branching for collision detection, and force relations (Fig. 1D-i; see Methods and Supplementary Fig. 3 for a full description).

We then instantiate dSPM’s two stages for this Ψ. First, for the dynamical algorithm, most of the board is simple to represent as a flow: between collisions, position integrates a constant velocity. The difficulty lies in two discrete, if-then events that a simulation-based representation requires: detecting collision (that the ball has reached a wall), and then reversing one component of its velocity and remembering this new direction thereafter. Neural populations, however, evolve smoothly. Following [35], we therefore build these events from a standard primitive of nonlinear dynamics, the bifurcation: a smooth change in one parameter that abruptly changes how many stable states a system has (Supplementary Fig. 4A). A common, textbook form, the pitchfork bifurcation, 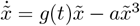, serves well for this purpose. While *g*(*t*) *<* 0, the system has a single stable state at 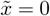 when *g*(*t*) rises above zero, 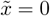 becomes unstable, and the state moves to one of two new stable states. Letting *g*(*t*) grow as the ball nears a wall turns 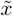 into a collision detector that stays at zero when the ball is far from the wall and switches on at contact. A second coupled cubic variable with two stable states acts as a memory element for the velocity sign, much like a flip-flop. It holds the sign of the velocity, is flipped by the collision detector, and keeps the new sign after the ball has moved away from the wall. This dependence on the history of collisions, rather than on the current position alone, is called hysteresis (Fig. 1E, phase-space graphs and board visualizations of each sub-panel; Supplementary Figs. 4A, 5). (The cubic form is not the only possibility; however, any implementation of collision-aware simulation needs, in some form, a bifurcation at the boundaries and a hysteretic memory of direction, and the cubic is the lowest-order polynomial that provides both. See also Discussion.) Coupling these primitives with linear integrators for position (Fig. 1D-ii, Supplementary Fig. 4B) fully specifies the algorithm in an M = 12-dimensional dynamical manifold (see Methods): much as a ring attractor captures the structure of heading direction, the resulting manifold maps the board’s physics.

Second, for neural falsifiability, we program this algorithm into the connectivity of a tanh-activated reservoir computer, a flow-matched embedding of 12 dynamical variables in *N* = 1, 000 units. To compare dSPM with neural data, for each Pong condition, we first initialize *r*(0) with that condition’s initial configuration 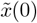 (the initial values of the 12 dynamical variables based on the ball’s initial position and velocity) by running the open-loop network (Eq. 1) for a small number of integrator time steps. We then unfurl the closed-loop programmed RNN (Eq. 3) starting from this “warmed-up” reservoir state (see Methods, Supplementary Information Section 1; Fig. 1D-iii). (In this instantiation, the dSPM does not receive any external inputs, thus 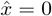 in both Eqs. 1 and 3.) These unfurled states of the reservoir’s “neurons” are what we compare to DMFC. Fig. 1E illustrates the resulting multilevel formulation (Supplementary Fig. 6, dynamic visualization). Crucially, during this initialization or at any stage of model development or testing, dSPM is never given or trained on any data, including DMFC data.

### Alternative methods: task-optimized RNNs and task-performant heuristics

To situate dSPM’s explanatory power in an appropriate context, we evaluate it alongside two alternative methods: (i) task-optimized RNNs and (ii) a task-performant heuristic strategy (a common explanatory framework in cognitive psychology [47]). We stress at the outset that this is not a model-comparison exercise among interchangeable candidates (e.g., as it is done when comparing deep neural networks with different architectures or capacities) — rather, the alternatives differ from dSPM in the kind of scientific inference they can, in principle, support about the brain. dSPM enables direct testing of algorithmic-level hypotheses — e.g., does DMFC predict by simulating the board’s physics and if so, what is that simulation mechanism like? Sufficiently expressive networks are universal function approximators [48, 49], and so the speculative possibility remains that a task-optimized network more powerful or different than the ones we are testing here could be crafted to match DMFC. However, finding one would still not reveal, on its own, the algorithm the system implements — instead, such inference requires additional post hoc arguments and interpretation techniques to extract the implicit, black-box procedures learned by the network [19]. (See also Discussion.) dSPM reverses the direction of inference. The algorithm is specified first, and the network’s dynamics are derived from it analytically, with a semiconjugacy guarantee; agreement or disagreement with DMFC therefore bears directly on the algorithmic-level hypothesis. In this design, the task-optimized RNNs serve as controls rather than as competing algorithmic hypotheses: they quantify what task performance alone induces in a recurrent network’s representations.

The first set of alternative models is the task-optimized RNNs, which solve the interception task, achieving near-ceiling performance by learning the statistical regularities needed to minimize their training objectives. One group of RNNs, which we call “Next-time point RNNs”, is trained on the combined objectives of predicting where the ball will be in the next time step and where to place the paddle for successful interception. Another group of RNNs, which we call “Paddle-only RNNs”, is trained only on the objective of where to place the paddle. These models are taken directly from refs. [26, 45], where their interception performance strongly correlated with the animals’ behavior in this task. We test both the GRU and LSTM variants of these RNNs and report pooled results due to their similarity.

The second alternative method probes DMFC with task-performant heuristics. Inspired by ref. [50], we evaluate dSPM against the ground-truth ball position and velocity (“Ball State”), which is shown to outperform large-scale video foundation models in representational similarity comparisons [50]. Ball State is a baseline in two ways. First, it quantifies the raw, stimulus-level similarity between Pong conditions, which is a similarity floor that any good account of DMFC must exceed. Second, because trajectories are piecewise linear in the interception task, the ball’s endpoint can be accurately linearly regressed from Ball State; this regression is a non-simulation-based strategy that predicts without representing physics, and one from which any algorithmic account of DMFC’s predictive ability must be distinguished. We now turn to the DMFC data.

### Rapid emergence of a low-dimensional, structured manifold in DMFC

The dSPM framework makes a structural prediction that precedes any comparison to neural data. Programming a physics-based representation of the board into the reservoir’s connectivity has a geometric consequence: the reservoir’s activity is confined to a low-dimensional manifold whose geometry mirrors that physics-based representation. No analogous structural constraint governs the task-optimized RNNs, whose hidden states are free to distribute representational content across whichever effective dimensionality minimizes their training objective. This asymmetry generates a testable prediction: if DMFC implements a physics-based representation of the board, its covariance structure should occupy the low-dimensional regime of dSPM’s programmed manifold rather than the higher-dimensional regime of the task-optimized alternatives. We tested this prediction through three complementary analyses of population geometry.

First, using principal component analysis, we computed the covariance eigenspectrum (the ranked percentage of variance explained by each principal component) for DMFC, dSPM, next time-point RNNs, and paddle-only RNNs, across all 79 conditions and time bins (Fig. 2A). The eigenspectrum of DMFC decays steeply, with variance concentrated in a small number of leading components and declining rapidly thereafter to the shuffled-data baseline (see Methods). The eigenspectrum of dSPM reservoirs follows a qualitatively similar pattern. As expected, there is more explainable variance in the model “neurons” than in the noisy measurements of the DMFC neurons, yet the variance decays to the shuffle-level baseline after a similarly small number of principal components. In contrast, next-time-point and paddle-only RNNs show substantially flatter eigenspectra, indicating that representational variance is distributed across many more dimensions.

**Figure 2.**
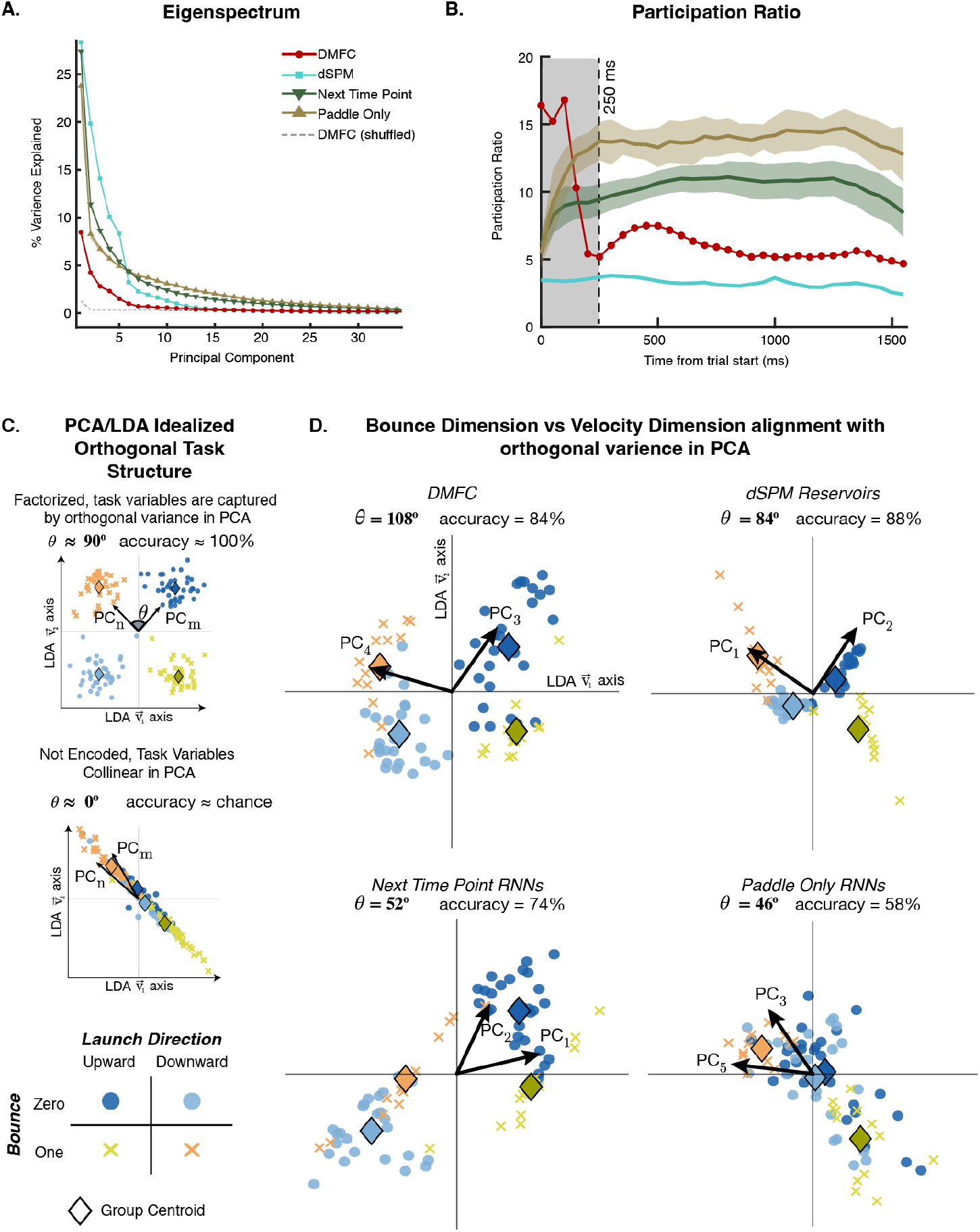
Latent Dimensionality and Neural Geometry in PCA Space. (A) Covariance eigenspectrum (percentage of variance explained per principal component) for DMFC, dSPM reservoirs, next time-point RNNs, and paddle-only RNNs, computed across all 79 conditions and time bins. DMFC and dSPM concentrate variance in a small number of leading components, decaying to the level of the shuffled-data baseline (dashed; DMFC with condition and time labels permuted, see Methods) by the 13th component, whereas the task-optimized RNNs distribute variance across many more dimensions. Shaded bands, SD across model instances; shuffle band, SD across permutations. (B) Time-resolved effective dimensionality (participation ratio) in 50 ms bins. DMFC (red) begins high-dimensional and collapses sharply at ∼250 ms (dashed line; gray shading, pre-transduction epoch), reaching toward the dSPM regime (cyan) and remaining well below the task-optimized RNNs for the rest of each condition. Bands, SD across model instances. (C) Logic of the geometry analysis, illustrated with idealized data. Four classes are defined by crossing the task’s two causal contrasts: bounce versus no-bounce, and launch direction 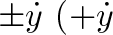, top wall; 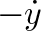, bottom wall). If the two contrasts are carried by orthogonal variance in the population’s principal components (top), the images of those components in the two-axis LDA discriminant plane remain orthogonal (θ ≈90^°^) and the four classes occupy separate quadrants with high classification accuracy. If the contrasts are collinear in the population variance (bottom), the imaged components collapse onto a single axis (θ ≈ 0^°^) and accuracy falls toward chance. (D) The same analysis applied to each system: all 79 conditions projected into the discriminant plane (v_1_, v_2_) at the best-discriminating principal-component pair and time bin (arrows, imaged principal components; diamonds, class centroids; four-way accuracy, chance 25%). DMFC (θ = 108^°^, accuracy 84%, PC_3_/PC_4_) and dSPM (θ = 84^°^, 88%, PC_1_/PC_2_) carry the two causal contrasts on near-orthogonal components, deviating from orthogonality by 18^°^and 6^°^. The task-optimized RNNs collapse toward oblique arrangements, deviating from orthogonality by 38^°^and 44^°^(next time-point, θ = 52^°^, 74%; paddle-only, θ = 46^°^, 58%).

Second, we quantified time-resolved effective dimensionality, i.e., the number of principal components actually needed to explain the variance in the data, using the participation ratio (PR) method [51, 52] (see Methods; Fig. 2B). DMFC’s effective dimensionality begins elevated at trial onset and undergoes a sharp, sustained collapse at ∼250 ms to nearly dSPM-level dimensionality. We note that PR is typically inflated by measurement noise and finite samples [53]. Because DMFC carries a noise floor that the deterministic models do not, both comparisons below are conservative: DMFC’s true dimensionality is likely closer still to dSPM’s, and its separation from the task-optimized RNNs likely wider. (Indeed, in Supplementary Fig. 7, we show that the gap between dSPM and DMFC is largely resolved after accounting for their different variability profiles.) The 250 ms collapse cannot be explained by sensory arrival alone. Visually evoked neuronal responses emerge earlier in the frontal cortex, e.g., with mean latencies of about 80-100 ms in the supplementary eye field, albeit for simple stimuli [54], placing the scene in the DMFC roughly 150 ms before the collapse. We attribute this residual interval to initial-condition inference: Unlike dSPM, which starts with its hidden state directly initialized for each condition (ball’s initial position and velocity; Fig. 1D-iii), so is low-dimensional from the start, the DMFC population must first estimate these initial board configurations for each condition. In the stable regime beyond 250 ms, both DMFC and dSPM show much lower effective dimensionality than the task-optimized RNNs.

Third, this dimensionality collapse raises a critical question: is the 250 ms manifold merely low-dimensional, or does its geometry encode something about the board’s physics-based structure? The 79 conditions offer two structural contrasts: whether the trajectory includes a wall collision (bounce versus no-bounce) and the launch direction (upward versus downward). In dSPM, these contrasts are carried by factorized (i.e., independent) dynamical variables (the bifurcation parameter for collision and the sign of the initial velocity), so the programmed manifold makes a specific geometric prediction that the values these two variables take across the 79 conditions will conform to two interpretable approximately ortgohonal axes.

To test this predicted geometry and compare it across systems, we performed time-resolved (every 50 ms from the start up to 500 ms, prior to any wall bounce) linear discriminant analysis (LDA) within the early-epoch principal component subspace (0–300 ms; Fig. 2C-D). For each system, we identified the 50-ms time bin and the pair of principal components that maximized cross-validated discrimination among the four classes of conditions defined by crossing the two structural contrasts (no-bounce upward, no-bounce downward, top-wall-bounce, and bottom-wall-bounce). Critically, this procedure does not maximize or select for any particular geometry: many different geometries can achieve the same accuracy. Yet, DMFC and dSPM not only achieved similar accuracy (84% and 88%; chance 25%), but also their geometries aligned, with the two discriminant axes near-orthogonal in both (*θ* = 108^°^and 84^°^, deviating from orthogonality by 18^°^and 6^°^). Both systems thereby orthogonalized the scene’s ‘causal degrees of freedom’, bounce versus no-bounce on one hand, and launch direction on the other: in the original PCA space, no-bounce conditions occupy a compact region along one axis, bounce conditions separate from them along an approximately orthogonal axis, and the upward and downward launch directions sit ∼ 180^°^apart along these axes. DMFC’s accuracy peaked in the 250-300 ms time bin, consistent with the collapse we observed in PR analysis, and this orthogonal geometry was observed across pairs of the second, third, and fourth principal components. In contrast, the task-optimized RNNs collapse toward oblique arrangements (*θ* = 52^°^and 46^°^, deviations of 38^°^and 44^°^), and their four-way accuracy is lower (74% and 58%). These results indicate that the observed DMFC geometry is not a consequence of just solving the task but is instead a signature of physics-based representation.

Together, these three analyses show that the DMFC recordings are consistent with a compact, physically-based manifold that emerges from the high-dimensional neural state space at the visual transduction latency and persists throughout the task, as dSPM predicts.

## Single-state sufficiency mechanism for rapid ball endpoint prediction

The dimensionality collapse and structured geometry established above motivate a mechanistic prediction: if the DMFC population state at ∼250 ms already encodes the physics-based structure of the scene in the orthogonalized geometry we observed, then a linear readout of that early state should be sufficient to predict future quantities, without any additional computation (Fig. 3A). In dSPM, this follows analytically from the manifold structure: each momentary hidden state encodes not only the current ball position and velocity but all 12 dynamical variables that determine the local curvature of the manifold. Because the local vector field of a dynamical system determines its global trajectories given sufficient initial constraints [55, 56], a manifold configured by perceived initial conditions should make future quantities available as linear projections along the surface of the manifold. This mechanism contrasts with step-by-step simulation in a physics engine (e.g., [3,26]) — the manifold geometry makes such simulation unnecessary.

**Figure 3.**
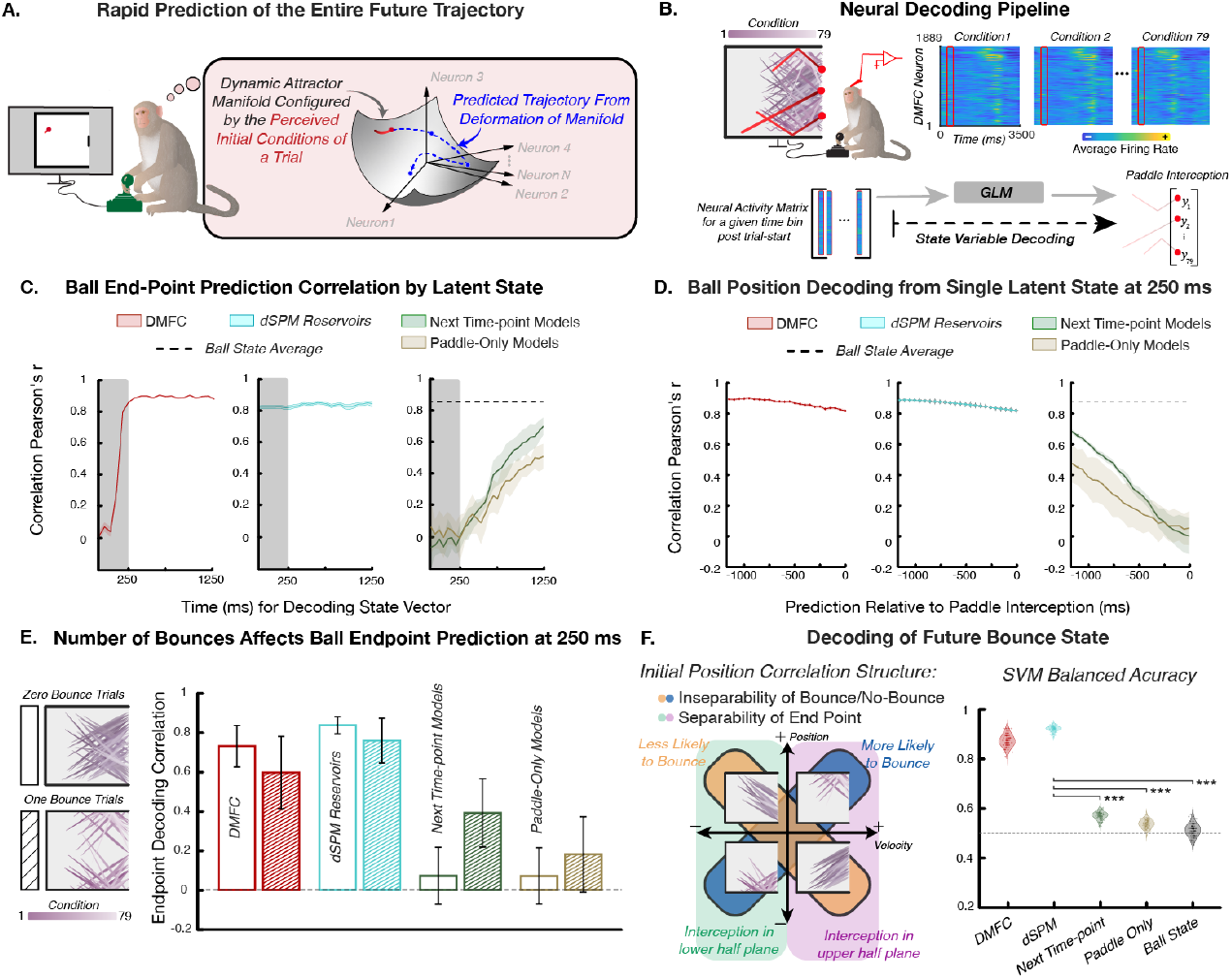
Confirmation of dSPM’s single-state sufficiency mechanism of physical prediction in DMFC activity. (A) dSPM posits that the perceived initial configurations of the gameboard bias the DMFC activity toward a low-dimensional manifold whose geometry corresponds to a physics-based representation of the gameboard. This manifold renders the board’s future trajectory lawfully decodable. (B) Our decoding pipeline is applied to both the neural data and models. This panel depicts the analysis of 1,889 DMFC neurons across 79 conditions in the dataset. For each time bin of 50 ms (from the start of the condition to the end), we used a generalized linear model (GLM) to decode the ball’s position (x-axis) at paddle interception (“ball endpoint”), and other aspects of the board state. (C) Temporal evolution of decoding accuracy for the ball endpoint from DMFC, dSPM, and task-optimized RNNs. DMFC neural activity (left, red) shows early prediction capacity ( ∼250ms after trial onset), with high correlation to actual ball endpoint (Pearson’s r > 0.8) maintained until late in the trial. This pattern is closely matched by dSPM (middle, cyan), while task-optimized RNNs (right, green/brown) show slowly improving prediction accuracy over time. (D) Decoding accuracy for the ball’s position across multiple points along its future trajectory (analyzed from −1450ms relative to paddle interception), using neural and model state vectors at 250 ms after trial onset. This confirms the striking prediction of dSPM: at the start, the complete future trajectory of the ball, not just the endpoint, is decodable in DMFC (Pearson’s r > 0.8 throughout for DMFC and dSPM), but not in task-optimized RNNs. Dashed lines in panels C and D are decoding performance using Ball State, which we further test in panel F. (E) Endpoint decoding at 250 ms, computed separately for zero-bounce (open bars) and one-bounce (hatched bars) conditions. DMFC and dSPM decode the endpoint robustly within both groups, with slightly better performance on zero-bounce conditions; the task-optimized RNNs decode poorly within either group. (F) Abstract bounce state representation prior to bounce. Left: bounce occurrence has XOR structure (i.e., linearly not separable) on the ball’s initial position × velocity, whereas endpoint decoding is a half-plane (i.e., linearly separable). Right: balanced accuracy of a linear SVM trained on pre-collision (250–500 ms) window-averaged states to classify held-out conditions as bounce vs. no-bounce; violins over cross-validation repetitions; dashed line, chance (0.5); stars, paired t-tests vs. dSPM. DMFC and dSPM classify far above chance; task-optimized RNNs and Ball State remain near chance.

We call this the ‘single-state sufficiency’ mechanism and first test it on the task-relevant interception point, the ball’s position at paddle contact. Rajalingham et al. [46] reported that the ball endpoint is decodable from DMFC remarkably early, at ∼250 ms after trial onset, and that no model they tested explained this rapid encoding. Our geometric analyses render this timing expected rather than mysterious. The participation-ratio collapse (Fig. 2B) independently identifies ∼250 ms as the moment DMFC’s population variance reorganizes, which dSPM interprets as the initialization of the condition’s manifold. It therefore predicts, from covariance structure alone and before any decoder is run, *when* future quantities should become linearly readable. The single-state sufficiency mechanism further explains why decodability arrives all at once rather than gradually accruing over time: once the manifold is configured, future quantities are already linear projections of the current state, and there is no simulation to wait for. To evaluate endpoint prediction across systems, we constructed a decoding pipeline (Fig. 3B) applying a generalized linear model (GLM) [57] to the neural or model state in each 50 ms time bin independently, predicting the ball’s endpoint position (see Methods).

Applying this pipeline to dSPM recapitulates DMFC’s rapid prediction with high fidelity (Fig. 3C, middle): decoded endpoints from dSPM’s hidden states track the neural pattern, maintaining strong correlation to the actual ball endpoint (Pearson’s *r >* 0.8) from early time points through each condition. Moreover, separating the 79 conditions into “zero-bounce” (direct trajectories without a wall collision) and “one-bounce” (trajectories with one wall collision) conditions (Fig. 3E), we find robust endpoint prediction within each group in both DMFC and dSPM, with slightly better performance on zero-bounce conditions.

The task-optimized RNNs, both next time-point and paddle-only variants, fail to reproduce this rapid endpoint prediction, replicating and extending Rajalingham et al. [46] (Fig. 3C, E), and paralleling their lack of dimensionality collapse and structured geometry (Fig. 2).

Rajalingham et al. [26] considered DMFC’s rapid endpoint decoding accuracy and the lack of such decoding in task-optimized RNNs as evidence for a distinct mechanism of simulation in the brain. dSPM provides such a mechanism through its physics-based manifold structure, which is absent in the task-optimized RNNs. However, accurate decoding does not, by itself, prescribe the computation underlying it [58, 59]. Indeed, for the current ball interception task, there is a competing non-simulation account that challenges this conclusion: the linear decoding of the ball endpoint from Ball State. The green and purple half-planes in Fig. 3F (left) show that endpoint interception in the upper versus lower half of the board is a linearly separable function of the ball’s initial position and velocity. Thus, simple linear regression on Ball State exploits this correlational structure of the task to achieve high-decoding accuracy without any simulation (Fig. 3C, right). We now turn to novel, structural predictions of dSPM that go beyond what can be explained with this task-performant heuristic.

## Confirming the entire future trajectory prediction of dSPM

The above analysis tests only a single, task-rewarded piece of information about the trajectory: the interception point. The dSPM model, however, makes a stronger prediction: because the ball endpoint occupies no special place in the physics-based manifold (it is simply one point among many on a globally structured trajectory surface), the manifold state at ∼250 ms should support linear decoding of any future ball position, not just the endpoint. Testing this prediction is therefore a direct probe of whether DMFC implements a physics-based representation, in which all trajectory information is simultaneously available from the early manifold state, or a task-efficient shortcut that encodes only the rewarded quantity.

We confirm this prediction in both DMFC and dSPM (Fig. 3D, left and middle). When decoding from the DMFC neural state at 250 ms, the ball’s position at any point along its future trajectory is reconstructed with high accuracy (Pearson’s *r >* 0.8), and the dSPM reservoir replicates this capability across the full trajectory. In contrast, task-optimized RNNs decode progressively worse at more distant future time points (Fig. 3D, right), consistent with a representational strategy tuned to the next state rather than the scene’s global physics. The agreement of each model’s decoding-vs-time curve with DMFC’s confirms this: dSPM’s concordance with the neural curve (CCC median 0.92, IQR 0.90–0.93, *n* = 50 reservoirs) far exceeds that of the task-optimized RNNs(0.03 and 0.01 for next-time point and paddle-only; exact rank-sum, Holm-adjusted *p* = 2.1 ×10^−9^, rank-biserial = 1.0); In this fine-grained analysis, dSPM also outperforms the Ball State heuristic despite its ability to exploit the correlational structure of the trials (0.90; one-sample *t* comparing dSPM reservoirs to Ball State level, *p <* .001; Supplementary Fig. 8). The entire future trajectory, not just the rewarded endpoint, is thus reconstructable from a single early population state, as the manifold geometry, and not the task objective, predicts.

To more stringently test dSPM’s physics-based representations, we focused on an aspect of the board state the task’s correlational structure cannot address: classification of a discrete future event whose dependence on the initial conditions is not linearly separable. Whether the ball will bounce, independently of the wall that it will, has a linearly inseparable (i.e., XOR) structure on the initial position and velocity: a ball high on the board bounces if moving upward but not downward; a ball low on the board, the reverse (the crossing yellow and blue bands in Fig. 3F, left). We trained a linear SVM on population states averaged over the 250–500 ms window, strictly before the earliest collision in the dataset (550 ms), to classify held-out conditions as bounce or no-bounce (balanced accuracy; chance = 0.5; see Methods). A linear classifier on the raw ball state cannot solve XOR, and Ball State accordingly performs at chance (balanced accuracy 0.51, AUC 0.48, *p* = 0.06), with the task-optimized RNNs faring only slightly better (balanced accuracy 0.57, *p* = 0.04, and balanced accuracy 0.54, *p* = 0.22, for next-time point and paddle-only). In DMFC and dSPM, by contrast, the abstract, wall-independent information about impending collision is overwhelmingly classifiable before it occurs (balanced accuracy 0.87, *p <* 0.02, and 0.91, *p <* 0.02; permutation test, 50 permutations). dSPM’s balanced accuracy exceeded that of Ball State, the next-time-point RNNs and the paddle-only RNNs by 0.39, 0.34 and 0.37, and that of DMFC by 0.03 (95% CI [0.027, 0.038]; all *p <* 0.001, direct bootstrap test on matched partitions; see Methods). These results provide further evidence for a dSPM-like physics-based manifold structure underlying DMFC activity, rather than task-optimized statistical representations or a task-specialized, performant heuristic.

### dSPM captures representational similarity of DMFC dynamics and subsumes what can be explained by alternatives

Finally, we turn to representational similarity analysis (RSA; see Methods), which imposes a qualitatively different test of the models [58, 60]. Unlike the decoding and geometric tests above, which target specific quantities or summary statistics, RSA evaluates the full pairwise similarity structure of population states across all 79 conditions and time bins simultaneously, and is therefore sensitive to the manifold’s nonlinear topology rather than its overall dimensionality or linear decodeability [58, 61]. Via partial correlation, RSA also tests whether dSPM retains explanatory power after accounting for the variance explained by the alternatives.

For a given system (either a model or DMFC), we built a representational similarity matrix, where each cell is the correlation between the activity at time-point *t*_*i*_ in condition *k*_*i*_ and the activity at time-point *t*_*j*_ in condition *k*_*j*_ (Fig. 4A). dSPM’s similarity matrix correlates substantially better with DMFC’s (*r* = 0.59) than do those of the next-time point RNN (*r* = 0.35), paddle-only RNN (*r* = 0.08), and Ball State (*r* = 0.32; all *p <* 0.001, direct bootstrap hypothesis testing against dSPM; Fig. 4B). Note that here, the Ball State alternative consists of the entire trajectory information, flattened vector of both positions and velocities throughout each condition (not just the initial position/velocity).

**Figure 4.**
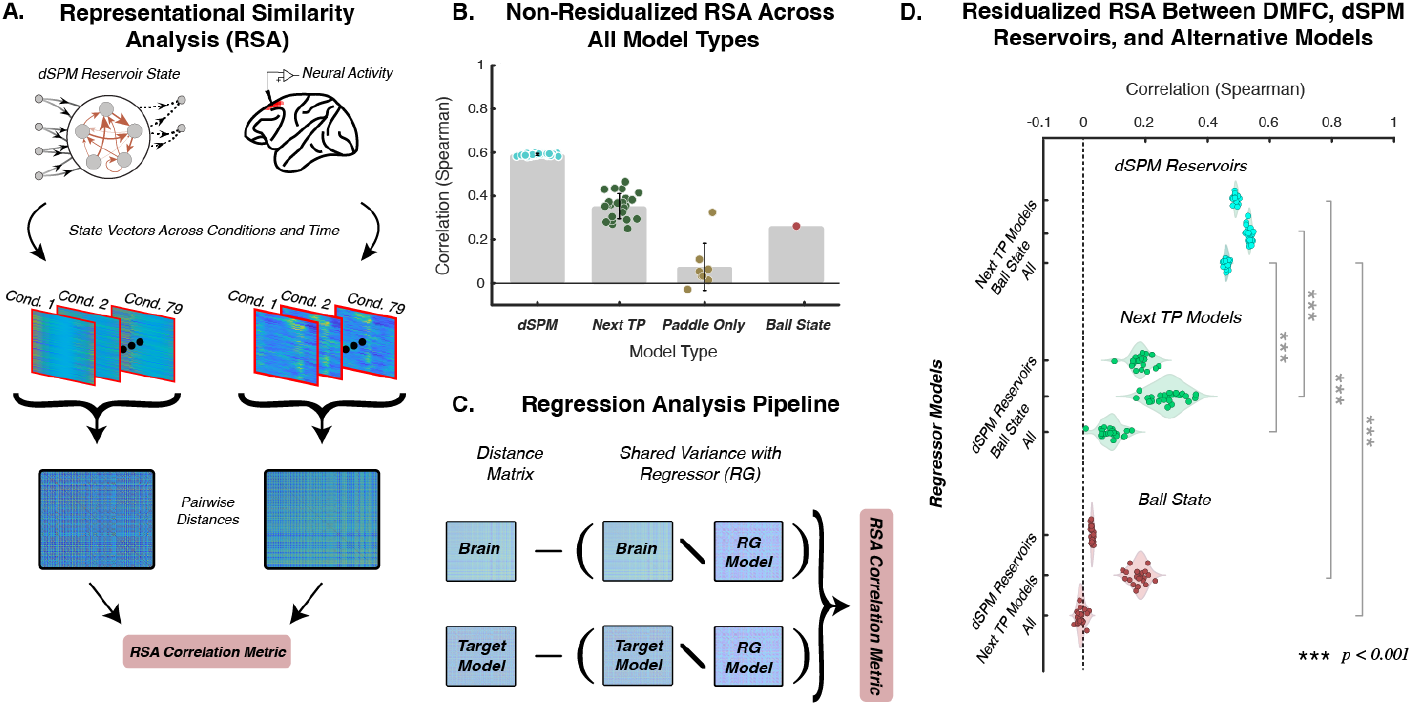
dSPM captures the similarity structure of DMFC neural dynamics across moments and conditions. (A) We performed a representational similarity analysis comparing the DMFC neural similarity matrix to the similarity matrices of each model. (B) dSPM reservoirs achieve a significantly higher correlation (Spearman’s ρ) than all alternatives. (C) A schematic of the partial regression pipeline to determine and compare unique variance explained by different model types. (D) Correlation and partial correlation (Spearman’s ρ) between model and neural representational similarity matrices are shown. The plot is organized by model type [top: dSPM; middle: next-time point; bottom: Ball State (based on flattened vector of all positions and velocities throughout a given condition)] and residualization condition (controlling for variance explained by different model types). Supplementary Fig. 9 shows the full set of residualization analyses. dSPM shows significantly higher correlations with neural data than alternatives across all residualization settings. Critically, when residualizing dSPM reservoir states from alternatives, there remains little explanatory power of these alternative models, while dSPM maintains a high correlation even after residualizing out all alternative models. This asymmetry indicates that dSPM reservoirs capture fundamental aspects of neural computation not present in alternative models.

We then used partial correlation to ask whether dSPM explains non-overlapping variance in the DMFC similarity matrix (Fig. 4C; see Methods). dSPM retains substantial explanatory power after residualizing both kinds of task-optimized RNNs; the next-time point model, by contrast, has almost no variance left to explain after residualizing dSPM and the paddle-only model (difference *p <* 0.001; Fig. 4D).

Ball State, as a measure of raw trajectory similarity, is a critical confound for any RSA over neural activity. Strikingly, dSPM not only outperforms this covariate (*r* = 0.59 vs. 0.32) but subsumes its entire share of the explained variance (Fig. 4D). In Supplementary Fig. 9, we provide a full residualization analysis across all pairs of models for an extended set of alternatives. Together, these results indicate a neural mechanism of physical prediction in DMFC: a latent, low-dimensional manifold whose geometry corresponds to a physics-based representation of the board, rather than task-optimized statistical representations or task-performant heuristics.

## Discussion

Distinctively, the present work synthesizes two prominent views of the brain, across the fields of cognitive science and neuroscience: The brain as a symbolic representation system of approximate structure-preserving maps of the world around us [6, 62] and the brain as a dynamical system [22]. Unlike typical cognitive models that focus on computational-level explanations and behavioral data, dSPM integrates across levels of analysis and offers falsifiable, distinctly interpretable hypotheses of neural mechanisms [63]. Instead of the popular data-driven approaches of interpreting low-dimensional manifolds in high-dimensional neural data, our approach provides a computationally constructive handle on these manifolds. dSPM joins a long lineage in theoretical neuroscience of starting from a stated computational objective and algorithm to infer properties at the implementational level, including sparse coding [64] and its neural implementations [65, 66], inference under linear forward models [67, 68], and more generally the class of computations that are specified as solutions of an optimization [32, 33]. However, dSPM qualitatively expands the set of possible computations to arbitrary nonlinear dynamical algorithms, including those that can run autonomously from an initial state without optimizing anything. Moreover, dSPM is distinct from low-rank connectivity inference approaches (e.g., [69–71]) in two ways. First, dSPM provides a direct handle at the algorithmic level (thereby connecting the computational-level objectives and implementation-level circuitry), whereas low-rank RNNs constrain the space primarily at the implementation level (as hypotheses over connectivity). Second, dSPM’s hypothesis space of computations is not constrained to what can be expressed with low-rank (typically linear) recurrent circuits; it can project any computational graph that can be expressed as a dynamical algorithm into an empirically falsifiable synthetic neural activity.

We implemented the dSPM framework by building on Kim & Bassett’s methodology, which provided a general-purpose tool for programming distributed recurrent computation and the general form of the Pong algorithm we applied here [35]. In doing so, dSPM creates a new, enabling bridge from applied math and physics to neuroscience and cognitive science — a previously unavailable multilevel modeling capability for identifying neural algorithms of complex mental representations. Commonly, manifolds in the brain have been explored by analyzing neural recordings with dimensionality reduction techniques, and formalized for simple settings tied to the animal itself (e.g., a ring for heading direction, a torus for position). In contrast, dSPM specified a manifold in advance for a physical scene out in the world (a substantially more complex domain close to those explored in cognitive science), and generated synthetic neural activity falsifiable in actual neural populations. This capability was not foreseen by [35] nor is it possible under the alternative modeling frameworks mentioned above. To facilitate adoption of dSPM, we provide an intuitive programming package to develop dSPMs without an extensive technical background (see Supplementary Section 2 and Supplementary Figs. 1-2, and Code and Data Availability).

With this new modeling capability at hand, this work suggested the ‘manifold-based simulation’ account of physical scene prediction in DMFC. It is the confluence of multiple independent lines of evidence that gives support for this account. dSPM predicts an early onset (identified at 250 ms in DMFC) of a manifold whose geometry corresponds to a physics-based representation of the scene, and four independent analyses confirm this: the collapse in dimensionality (Fig. 2B), the orthogonal separation of task variables in the data-driven latent space (Fig. 2D), the onset of endpoint and full trajectory decoding (Fig. 3C, D), and the binary classification of the task’s key nonlinearity before it happens (Fig. 3F). Moreover, dSPM continues to explain DMFC’s representational similarity of the task conditions after removing all the variance that can be explained by alternatives, whereas it subsumes nearly the entire explanatory power of the same alternatives (Fig. 4D). Although different alternatives, to varying degrees, are consistent with subsets of this range of phenomena, it is only the dSPM that not only anticipates these phenomena but also empirically demonstrates the alignment of its predictions with the neural data. That is our evidence that this frontal population is organized by a physics-based representation of the scene, and not by the demands of the task or the statistics of its input alone.

We interpret the rapid collapse in effective dimensionality of the DMFC population as the geometric signature of manifold initialization (Fig. 2B): perceived initial conditions of a scene rapidly collapse high-dimensional neural activity onto a low-dimensional manifold whose geometry corresponds to a physics-based representation of the encountered scene. This result can be seen as an example of the more general phenomenon that stimulus onset quenches neural variability [72]. Thus, computationally capturing this sensory-input-driven rapid collapse with dSPM could provide a concrete mechanism for this widely observed phenomenon of the central nervous system. Under dSPM, we can interpret this collapse as the Bayesian transport of a broad a priori possible range of manifolds to a ‘posterior manifold’ conditioned on the sensory inputs. Extending dSPM with such an inference process is of high priority.

Manifold-based simulation resolves a puzzle of intuitive physics identified in [26], explaining why and how a far out future event is encoded in neural population activity. In so doing, it simultaneously achieves two computational goals that have typically been considered distinct (e.g., [26, 73]): rapid, long-time-step predictions of future trajectory, available immediately upon manifold initialization as a linear projection along the manifold’s surface, and ongoing local tracking of the ball’s current position, available continuously as a local readout of the manifold’s tangent direction at each moment. Crucially, these are not two separate computations running in parallel but a single geometric object with both properties: the global geometry of the manifold implies the local tangent direction at every point, and the local tangent direction is the next-step prediction. The brain thus achieves what might otherwise require two dedicated systems — a rapid offline predictor and an online next-step tracker — through a single physics-based manifold representation that provides both simultaneously. We also note that this mechanism seems consistent with the phenomenology of playing a game like Pong: when observing a ball trajectory, we can make far-out predictions (e.g., what and where the ball will collide next) almost immediately in one’s mental representation, while also simultaneously tracking the moment-to-moment actual position of the ball.

This computational mechanism also has implications for the broader architecture of perception and planning and reasoning in the brain. Computational formulations of perceptual processing require moment-by-moment prediction, whereas planning computations can afford larger time steps through a more qualitative form of prediction. It remains unclear how these two kinds of predictions — moment-by-moment versus longer-horizon, more qualitative predictions — relate to each other, computationally and neurally. Manifold-based simulation suggests an exciting possibility. The relevant “error signal” it can support includes not only “the ball is not where I predicted it to be right now” but also “the ball’s trajectory is inconsistent with the manifold I initialized”. This latter type of manifold-level error signal would be capable of detecting not just momentary positional discrepancies but violations of the underlying physics — a ball that unexpectedly curves, accelerates, or passes through a wall would generate an error not because its current position differs from a point prediction, but because its trajectory is inconsistent with the manifold’s topology. In this way, the manifold-based simulation mechanism offers a path toward an integrated architecture of cognition, in which perception and planning/reasoning operate over substantially overlapping computational substrates. Future work should develop these integrated computational architectures with a manifold-based simulation core, and neurally test them in tasks that involve tight coupling among perception, physical prediction, and planning (e.g., the predator-pursuit task in [74]).

It is important to distinguish between the level at which dSPM makes claims about DMFC. At the algorithm level, our claim is not that DMFC uses the specific differential equations that make up dSPM’s dynamical algorithm, but that it implements the same equivalence class as dSPM: an algorithm capable of collision-aware physical simulation with hysteretic velocity memory requires, in some form, bifurcation at collision boundaries, hysteretic state maintenance, and differential time scales between these components. The eigenspectrum, participation ratio, geometry, decoding, and RSA analyses all operate at this equivalence-class level. Similarly, at the implementation level, the dSPM reservoir is a rate-coded network, with comparisons made to trial-averaged firing rates in brief time bins, where DMFC neurons spike; future spiking extensions of dSPM reservoirs are of pressing interest.

How could dSPM-like manifolds be learned from experience? Our results provide three threads of guidance. First, for the model class of task-optimized deep neural networks (where the researchers primarily have control over the training datasets, training objective, architecture, and sometimes the learning rule), our comparisons with task-optimized RNNs of Rajalingham et al. [26, 75] establish that it is not sufficient to only train for task performance or next-time-point prediction. Future work can consider finer-grained supervision when training such networks, as explored in the machine learning for learning dynamical systems (e.g., [76, 77]). Second, another intuitive possibility is video foundation models trained on much larger real-world datasets (e.g., [78–80]). The present study provides evidence that this recipe, at least by itself, may not be a “silver bullet”: The Ball State covariate we tested here is the same as the “Oracle” in ref. [50], which showed that it outperformed all of such large-scale video models they considered. Third, a core aspect of dSPM is the physics-based factorization of scene elements that underlie the scene — and this factorization is evident in the geometry of the dSPM reservoir as well as DMFC. Thus, we suggest that explorations of learning dSPM-like representations should incorporate such an ability to factorize the scene, either through their architecture, training data, training object, or their combinations. We once again emphasize that dSPM provides a uniquely direct and powerful handle at the algorithmic level. Task-optimized RNNs primarily constrain the solution space at the computational level (with their objective functions and training dataset), and to some degree also at the implementation level due to architecture. dSPM, in contrast, allows direct specification of algorithms, connecting computational-level objectives and implementation-level circuitry in a principled fashion. As a boundary condition, consider training a traditional RNN to predict not just the next time point and not just the ball’s position (as in the next time point RNNs), but all 12 variables of dSPM’s dynamical algorithm for the whole future trajectory. In the terminology of Sagodi & Park [81], the output of such a network would achieve high ‘similarity’ with dSPM’s 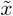, but it may do so while incurring a ‘complexity’ penalty, deforming the flow, 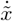. In contrast, dSPM’s reservoir, by construction, embeds the flow (Eq. 5, Methods). To the extent that one wishes to explore well-defined algorithmic hypotheses, dSPM offers a unique vehicle.

The current study motivates us, and hopefully other researchers too, to pursue extensions of dSPM to other domains of cognition, including representations of agents (e.g., [82]), places [83], and more complex physical scenarios (e.g., [5]). The key to achieving this generalization is to build dynamical algorithms for target domains and their computational graphs. An exciting future direction is to make the development of dynamical algorithms for reservoir computers systematic, much as high-level programming languages did for conventional computer architectures. The broad knowledge base of dynamical algorithms (e.g., [35–38]), currently dispersed across different fields, provides starting points for such a language. With such generalization and modeling capability at hand, dSPM could serve as a general tool for testing hypotheses about the computational foundations of biological intelligence, and for connecting algorithmic-level hypotheses to how high-dimensional recurrent circuits can actually realize them.

## Methods

### Dynamical Structure-Preserving Manifolds (dSPM)

Here we provide the details of the computational, algorithmic, and implementation levels of dSPM.

#### Computational-Level Description: Structure-preserving, physics-based representations for physical prediction

At the computational level, physical prediction can be described as building and manipulating physics-based representations of the physical world [84], flexibly supporting downstream adaptive behaviors. Following Battaglia et al. [84], we provide a structure-preserving physics-based representation of the gameboard in Supplementary Fig. 4. This symbolic program describes the entities (objects, walls, paddle) and their dynamics and interactions, using a high-level object-oriented programming language. The dSPM framework transforms such computational-level hypotheses, which lack neural grounding, into falsifiable proposals of neural mechanisms.

#### Algorithmic-Level: Dynamical Algorithm

Recent work in neuroscience provides evidence that basic structure-preserving representations, such as one’s heading direction or 2D position in space, are encoded in low-dimensional, latent manifolds underlying high-dimensional neural activity. These manifolds contain attractor dynamics (stable and unstable points of attraction and repulsion) that functionally map entities and their relations in the world. To realize complex cognitive representations, we need to express significantly more sophisticated computations in these dynamics.

To do so, we build on Kim & Bassett [35] to formulate a dynamical algorithm of a physics-based representation of the gameboard. This dynamical algorithm (Supplementary Fig. 4, 5) is a set of 12 coupled differential equations whose outputs are denoted as 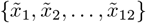. This dynamical algorithm defines a dynamical manifold, meaning that it’s a manifold whose attractors and repellers change over time. These dynamical attractors — i.e., nullclines in phase space that change with time *t* — correspond to a physics-based representation of the gameboard, including position updates, as well as collision detection and resolution. The 12 dynamical variables implement four main computational blocks (corresponding to the four code blocks in Supplementary Fig. 4) based on two forms of third-order polynomial logic.

We first describe these polynomials.

- **Pitchfork bifurcation** where *f*(*t*) controls whether we have a single stable attractor at the origin, (*f*(*t*) 0), or two symmetric non-zero stable fixed points at 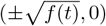 and one unstable attractor at the origin, (*f*(*t*) *>* 0), (textbook example, [85])

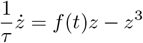
- **Polynomial-Shift Operator** The second cubic simply slides the attracting point left or right without changing its stability. The time-varying term *f*(*t*) raises or lowers the cubic, while the constants *α* and *β* tune the slope so that |*z*| remains ≤ 1. Keeping the state in this range guarantees that the activity vector of our recurrent network never leaves the ball-park set by the weight matrix’s spectral radius (roughly, the largest eigenvalue magnitude) [42], preventing runaway excitation

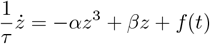

In these equations, the coefficient *τ* controls the sensitivity of the dynamics encoded in these polynomials to local changes (as well as initial conditions). Specifically, when |*τ*| ≫0, then the derivatives on either side of the fixed points are much stronger (Supplementary Fig. 5), making slight changes in the system more sensitive to changes. (This time constant is multiplicative to the global time constant of our reservoir computer, represented by *γ* in our update equation in the next section.)

The four computational blocks of our dynamical algorithm are as follows.

1. Constant registers (velocity components) To make each trial’s initial velocity available to the rest of the network, we reserve two dedicated state variables, *z*_1_ and *z*_2_, whose dynamics are purely integrative registers:

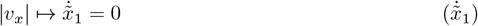

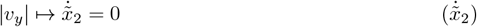

Because 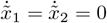, these variables act as constants during the simulation, giving the dSPM a read-only handle on (*v*_horizontal_, *v*_vertical_) while re-using the same recurrent weights across all trials.
2. Velocity Logic

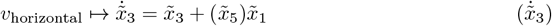

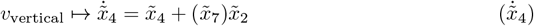
3. Hysteretic Switch for Nonlinear Reflections. Each state variable takes values in [−*x*_*c*_, *x*_*c*_], with

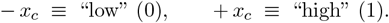

The dynamic logic couples two recurrent variables, the wall-collision variable 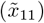 and the opposite hysteretic variable 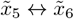 and 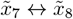

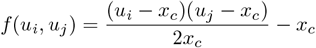

implements a Boolean NAND gate:

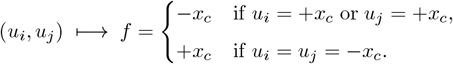

Because the output itself sits at a stable fixed point ( ±*x*_*c*_), it retains its state against small perturbations — i.e. it forms a hysteretic memory element. Cross-coupling two such units yields a bistable attractor manifold that we use as the circuit’s nonlinear reflection (bounce) switch.

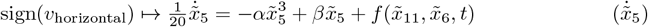

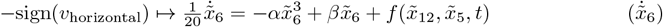

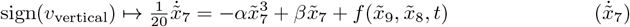

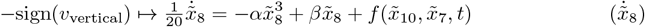
4. Collision Detection, where the extent of the board is defined by its height (*h*) and its width (*w*), *p*_horizontal_ and *p*_vertical_ are the paddle’s horizontal position and vertical position respectively, and *ϵ* and *σ* define the sensitivity/resolution of the collision detector relative to the ball’s distance to the walls/paddle.

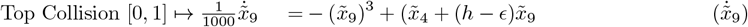

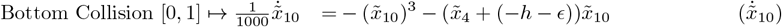

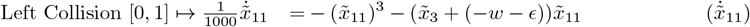

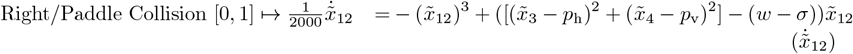

### Implementation Level: A Reservoir Computer Analytically Embedding the Dynamical Algorithm

In contrast to the traditional task-optimization approach that learns the weights via numerical training in a deep neural network, we constructed a separate class of models, i.e., reservoir computers, by analytically embedding the dynamical algorithm defined above (the twelve differential equations encoding ball motion, wall collision logic, and velocity sign changes) into the reservoir computer’s connectivity weights. We do so by following the method of Kim & Bassett [35].

The basic steps of this method are as follows.

1. **Start from the open-loop reservoir, a continuous-time network of leaky integrators:**

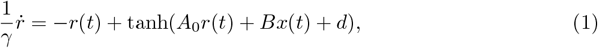

with tanh nonlinearity. We draw *B* i.i.d. from *U*[ −0.5, 0.5].Here *x*(*t*) is the full input vector and *B* the full input matrix. When the loop is closed (Step 6), this term splits as 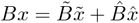, where 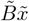 carries the inputs that will be programmed and fed back as recurrence, and 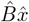 carries any external drive to the closed-loop system. Because the open-loop programming in Steps 2 through 5 treats all inputs symbolically and identically, the split is immaterial there, and we work with the combined term *Bx* until the loop is closed. For Pong there is no external drive 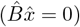; we retain the term for generality. *A*_0_ is the initial connectivity, which in the Pong setup is set to 0 for the open-loop solution and will be folded into the “programmed” connectivity in the closed-loop network. (Note that there are multiple notational conventions for expressing the time constant of a differential equation: 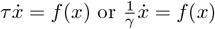. Here we use 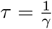, in line with the convention of Kim & Bassett [35].)
2. **Solve hidden state as a function of inputs**. We choose a random operating point, *r*^*^, and set the bias so that *r*^*^ is the resting state of the open-loop network at input *x*^*^:

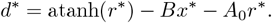

In open loop, each unit is a leaky integrator driven by a nonlinear function of the inputs, so the solution of Eq. 1 is determined by the inputs and their time derivatives and can be expressed as a smooth map

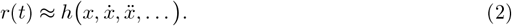

This map is used only to derive the connectivity; the closed-loop network that is run is fully nonlinear.
3. **Symbolic expansion**. Expand Eq. 2 to *k*^th^ order via a multivariate Taylor series, obtaining a “design matrix”

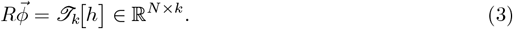

Each column is a monomial basis function represented by the symbolic vector 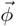 of the inputs, their time derivatives, and the multivariate interaction terms up to order *k*.
4. **Program the desired vector field**. Let *P* ∈ R^*M ×k*^ hold the same monomials, 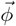 but evaluated on the *target* dynamics 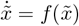. Compile the programmed readout

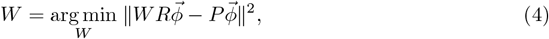

where ∥ · ∥^2^ denotes the Frobenius/*L*_2_ norm. This yields 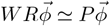 and hence

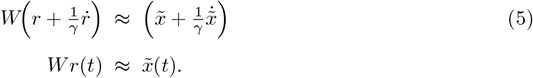
5. **Load initial conditions**. Each trial is specified by its starting state 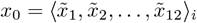: the ball’s speed along each axis, its position, the state of each direction switch (set by the sign of the velocity), and the four collision detectors at rest. With this input held fixed, the open-loop reservoir is integrated from *r*^*^ until it settles at its *conditioned operating point*, 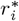:

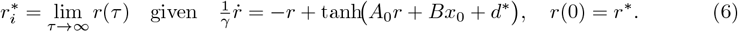

The integration uses Runge–Kutta 4 (RK4), and the reservoir converges within a few integrator steps. The symbolic initial conditions, *x*_0_, thus act like a key: they select, in the reservoir’s high-dimensional state space, the point that corresponds to that trial’s initial conditions in the low-dimensional target system. This “trial-conditioned” state, 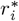, is the analytic operating point *r*^*^ displaced by the input *x*_0_, and it is the initial condition for the closed-loop reservoir (Step 6).
6. **Close the loop**. Split *Bx* into a recurrent part 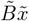 and an external part 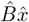 (this term is zero for Pong because there is no external drive), then substitute 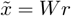:

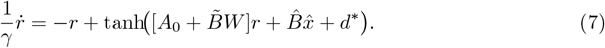

This yields the programmed adjacency 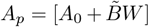. From 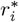 onward, we simply integrate the closed-loop reservoir (Eq. 7) with RK4. Nothing in this update equation refers to a ball, a wall, or a collision: the reservoir evolves only according to its own connectivity. The board dynamics are latent in that evolution and become observable only through the linear map

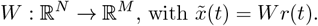

Released from 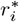, this readout traces the trajectory that the dynamical algorithm would take from *x*_0_. This is what Steps 1 through 5 work toward. The linear readout, *W*, matches the reservoir’s flow-field expansion, 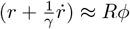, to that of the target, 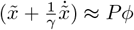, making the two systems semiconjugate in their phase-space topology, to the order of the expansion (Step 3); in particular, the fixed points of the target map onto fixed points of the reservoir. This semiconjugacy is what links the levels of the framework, translating between the algorithmic level of the symbolic equations and the implementation level of the recurrent network of leaky integrators.

Full derivations and hyperparameter choices appear in Supplementary Material “Details of Programming Reservoir Computer”.

#### Spectral radius of the programmed adjacency

In conventional reservoir computing, the adjacency matrix is drawn at random and rescaled so that its spectral radius, the largest magnitude among its eigenvalues, is below 1 [25, 42]. This is the criterion conventionally used to ensure the echo state property: with coupling this weak, the reservoir’s activity decays toward the origin, 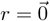, in the absence of input, and only a linear readout is trained. The programmed adjacency of Step 6 is not like that. The adjacency matrix *A*_*p*_ is solved/programmed, not drawn; and it is not programmed about the origin, but instead, it is programmed about *r*^*^, which typically has a large magnitude. For the Pong reservoirs, the spectral radius is large, *ρ*(*A*_*p*_) = 4.1 *×* 10^3^ *±* 0.4 *×* 10^3^ (mean *±* s.d. across 50 reservoirs; range 3.4 to 5.0 *×* 10^3^); by the conventional measure, these are strongly coupled networks far outside the echo-state regime. This value of the spectral radius, however, treats *A*_*p*_ as though the dSPM reservoir operated about the origin. See Supplementary Section 3 (and Supplementary Fig. 3) that accounts for dSPM’s non-zero operating point and a time-resolved analysis of its spectral radius, illustrating how the *>* 1 regime, avoided in traditional reservoir computing [25, 86], is instead a controlled part of the computation in dSPM.

### Applying dSPM to Experimental Conditions

Once we have our programmed reservoir computer, we can “load in” the initial conditions (Step 5) and evolve our network over time (Step 6). As the reservoir evolves its distributed computation, across the *N* hidden units, together computes the dynamical algorithm that constitutes our physics-based representation of the gameboard.

We initialize dSPM with the initial conditions used within the monkey experiments. The initial conditions contain ball position and velocity for each trial, which dSPM evolves autonomously without external inputs for the duration of the given Pong condition.

### Alternative Models

Rajalingham et al. [45] trained a large ensemble of standard machine-learning-style RNNs on the same interception task, using supervised learning protocols. These networks — here referred to as “task-optimized RNNs” — took the ball’s visual inputs and were optimized to predict and/or control the paddle’s position in order to intercept the ball.

The next time-point models were trained to predict both the ball position in the next time step and to control the paddle (where the paddle should be for successful ball-interception at the end of the trial). They considered different variants of the next time-point models depending on the specifics of the loss and architecture, which we pool together as they did not differ from each other statistically. The paddle-only models were trained only the latter objective — the ball endpoint.

They considered different hyperparameter choices [e.g., number of units, RNN circuit type (GRU, LSTM), input representation, etc.]. Here we report the best performing variants (number of hidden units=40; RNN circuit type=LSTM or GRU, input representation=motion filters; pixel input or Gabor-filtered input). We used these task-optimized RNNs “as is”, extracting their hidden states on each condition for comparison with DMFC recordings and dSPM.

### Stimuli and Neural Data

Two adult macaque monkeys (*Macaca mulatta*), one male (Monkey P) and one female (Monkey M), participated in this study. Animals performed a naturalistic ball interception task developed by Rajalingham et al. [45, 46]. In this task, each trial began with the ball in a random initial position and velocity in a two-dimensional arena presented on a digital monitor. The ball traveled rightward at a constant speed, with zero or one bounce off the horizontal walls. Crucially, the ball was rendered only for the early portion of each trial; its trajectory then became occluded before reaching the far right side. The monkeys controlled a paddle positioned at the right edge via a one-degree-of-freedom joystick, attempting to intercept the ball upon its (unseen) arrival. Each trial ended when the ball either made contact with the paddle or exited the right boundary of the display. Inter-trial intervals were 750 ms. Monkeys were rewarded with a juice drop when they successfully intercepted the ball. They were free to move their eyes during the occluded period and routinely shifted gaze in ways consistent with anticipating future ball positions.

Neural signals were recorded from DMFC using high-channel-count silicon probes (Neuropixels) in one animal (Monkey M) and linear probes (Plexon V-probes) in the other (Monkey P). Recording sites spanned an 8 mm × 3 mm grid in Monkey P (24 distinct locations, each sampled in two sessions) and a narrower grid in Monkey M (6 locations, each sampled in one session). Neurons were neither preselected nor excluded based on their response properties, and recording sites were not chosen based on putative task selectivity. Spikes were sorted using an automated algorithm (Kilosort 3.0) and subjected to quality checks that removed unstable or noisy units. Per-trial, per-neuron spike counts were binned in 50-ms intervals. Only units with significant split-half reliability (*p <* 0.01) were retained, resulting in a final dataset of 1,889 reliably recorded neurons across both monkeys. Trials were pooled into 79 unique stimulus conditions by averaging spike counts across repetitions of each condition.

### Neural Data Analysis and Model-Data Comparisons

#### Eigenspectrum Analysis and Participation Ratio

For the eigenspectrum analysis, for each system (DMFC or a given model), we constructed an input matrix of dimensions [*C ×N*] where *C* flattens the 79 conditions and the time bins each condition lasts, with 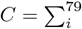 length(condition_*i*_), and *N* is the number of units (e.g., 1,889 units for DMFC, 1,000 units for dSPM, etc.). We then computed the covariance matrix of this input matrix and extracted its eigenvalues; the eigenspectrum is the ranked percentage of total variance explained by each component. A shuffled baseline eigenspectrum of the neural data was computed by applying a global element-wise permutation of the entire input matrix (following ref. [26]), destroying all temporal, cross-neuron, and cross-condition structure while preserving the marginal distribution of firing rates. The shuffle was repeated 20 times, and the mean and standard deviation of the shuffled eigenspectrum are reported.

For the PR analysis, the input matrix is structured differently. Because this is a time-resolved analysis, we cannot include time bins for which there aren’t sufficiently many conditions. As a trade-off between the amount of data and meaningful analysis, we set a cut-off of 32 time bins (1550 ms) where 90% of the conditions are still active. (Thus, the input matrix is of dimensions [*C ×N*] where *C* = 79 × 32.) Using this input matrix, we performed a global PCA, computing its covariance matrix and extracting the eigenvalues for the top 100 principal components. We then computed PR at each time bin *t* using these components:

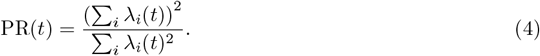

where *λ*_*i*_(*t*) is the eigenvalue of the *i*^*th*^ principal component. This equation provides an estimate of the actual dimensionality underlying the data for each time bin. [52, 87]

For the early-epoch PC geometry analysis, for each system, we restricted the input matrix to the 0–300-ms period (i.e., the first 6 time bins) and computed the covariance matrix’s eigenvectors corresponding to the top 15 components. We then projected each state vector of this system for each time bin (from time bin 1 to 32) onto these early epoch-specific eigenvectors. Over pairs of eigenvectors, linear discriminant analysis (LDA) was performed to identify the time bin and PC pair that led to maximum between-class separability for the three bounce-type categories (no-bounce, top-bounce, bottom-bounce); the cosine angle between the discriminant axes and the discrimination accuracy are reported as measures of orthogonalization.

#### Geometry of bounce and initial velocity direction

For each system, we asked how two structural contrasts of the task are arranged in the population state: whether the ball will bounce off a wall before reaching the paddle (bounce versus no-bounce), and the sign of its initial vertical velocity (launch direction, upward versus downward). Crossing the two contrasts defines four classes of conditions: no-bounce upward, no-bounce downward, top-wall bounce and bottom-wall bounce. Principal component analysis was performed separately for each system on its states restricted to the early epoch of the trial (0–300 ms, the first six 50 ms bins), and the first 15 components were retained. The states of all conditions at every time bin were then projected onto these early-epoch components. For every pair of components, and every 50 ms bin up to 500 ms (before the earliest bounce at 550 ms), we computed two discriminant axes in the plane of that pair, each the difference between the mean states of two groups of conditions, normalized to unit length. The first, 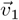, separates upward-from downward-launched conditions. The second, 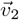, carries the bounce contrast: because a bounce reverses the ball’s vertical velocity, whether a condition bounces is fixed, given its launch direction, by the direction from which the ball arrives at the paddle, and 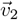 separates upward-from downward-arriving conditions. These are linear discriminant axes under an isotropic within-class covariance. A condition was scored as correct if its projections onto 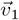 and 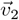 had the signs of its class, and for each system we selected the pair of components and the time bin with the highest four-way accuracy over all 79 conditions. We report this accuracy and the angle *θ* between 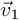 and 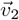 at the selected pair and bin. Because the sign of each axis is fixed by its definition (upward minus downward), *θ* is measured without folding, over the range [0^°^, 180^°^], with 90^°^indicating orthogonal axes. For Ball State, the four raw kinematic variables were used in place of principal components.

#### Trajectory Decoding from State Vectors

To test the future decoding capabilities of latent state representations of DMFC and different models, we used generalized linear model (GLM) [57] fits to decode the ball’s position from the latent state vectors over time. Neural population responses, task-optimized RNN hidden states, linear map, and dSPM reservoir states were all compared via two complementary analyses: ball endpoint decoding and full trajectory decoding. All GLM results reported are cross-validated using nested 5-fold cross-validation with 4-fold inner cross-validation for hyperparameter selection.

For ball endpoint decoding, we regressed each population’s activity at each time point onto the ball’s actual position at the time it would leave the board (i.e., where it would be intercepted by the paddle). For full trajectory decoding, we regressed each population’s activity at 250 milliseconds onto the ball’s actual position along its trajectory (including endpoint). The *x*-axis in Fig. 2(c,d) represents time relative to paddle interception (referenced as 0 ms). Since trials varied in duration due to different ball velocities and starting positions, we aligned all trials to their endpoints. To have consistent training/testing splits across all time bins, we could only decode positions going back 1450 ms from interception, corresponding to the duration of the shortest trial (29 time bins × 50 ms).

#### Classification of an upcoming bounce

We asked whether each system’s population state, before any bounce has occurred, linearly separates the 26 conditions in which the ball will bounce from the 53 in which it will not. For each condition, the state was averaged over the 50 ms bins from 250 to 500 ms (the earliest bounce occurs at 550 ms), giving one feature vector per condition. A linear support vector machine was trained and tested with 5-fold cross-validation over conditions, with features standardized on the training folds. The regularization constant was chosen within each training fold from {0.01, 0.1, 1, 10, 100} by an inner 3-fold cross-validation. Predictions were pooled across held-out folds and scored by balanced accuracy, the mean of the hit rates on the two classes, for which chance is 0.5 despite the unequal class sizes. This procedure was repeated over 100 random partitions, the same partitions for every system; for systems with multiple instances, balanced accuracy was averaged over instances within each partition, and we report the mean over partitions. A null distribution was obtained by permuting the bounce labels across conditions (50 permutations, regularization constant fixed at 1). Differences from dSPM were assessed by direct bootstrap hypothesis testing on the matched partitions: the paired differences in balanced accuracy were resampled with replacement 1,000 times, and *p* is the fraction of bootstrap samples in which the mean difference did not favor dSPM; the 95% confidence interval is the percentile interval of the same bootstrap distribution.

#### Representational Similarity Analysis

Representational similarity analysis (RSA) evaluated the topology of each system’s representations (model or DMFC) by computing pairwise Euclidean distances between its states for each pair of conditions and time bins, yielding a representational dissimilarity matrix (RDM) for each system of dimensionality [*C* ×*C*] where 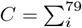 length(condition_*i*_). We then computed Spearman correlations between the upper triangular portions of the model and neural RDMs to assess their representational similarity. To determine whether dSPM captured unique variance in neural representations beyond task-optimized models and linear map, we employed a residualization procedure. For each model type, we regressed out the contribution of alternative models from both the neural and model (upper triangular, flattened) dissimilarity matrices using ordinary least squares:

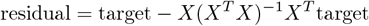

where *X*^*T*^ *X* contains the regressor dissimilarity matrix. Residualization was performed as follows: We randomly sampled a single model instance from the source model class as the regressor, then repeated this procedure across all model instances to account for sampling variability. This procedure was applied bidirectionally, residualizing dSPM models from task-optimized models and vice versa. Because the linear map heuristic has no variability (as it is based on regressing from the ground truth position and velocity information), our method carries over the variability due to dSPM instances for statistical quantification. We report all possible residualization pairings in supplementary figure **??**. Statistical significance was determined using parametric pairwise comparisons between residualized correlation values (paired samples two-tailed t-test). The asymmetric pattern of residualization results — where dSPM maintained high correlation with neural data after removing task-optimized and linear map’s variance, but not vice versa—indicates that dSPM captures fundamental aspects of neural computation not present in the alternative models.

#### Computational Implementation

All analyses were performed in MATLAB. Where parallelization was beneficial, data splits and model simulations were distributed across a high-performance cluster, with each trial or cross-validation fold assigned to a separate worker. The same 79 task conditions were used in both neural and model analyses, and all time series were binned at 50 ms. Optimal hyperparameters in regression or decoding analyses (e.g., ridge penalty terms) were chosen via nested cross-validation. The final outputs (hidden states, regression weights, decoded trajectories) were stored and evaluated with identical metrics for both neural and model data.

## Code and Data Availability

All methods for data preprocessing, model simulations, and decoding analyses were implemented in MATLAB and will be made available through a public repository.

## Acknowledgements

We are grateful to Mehrdad Jazayeri for discussions about this project and generously sharing neural data. We also thank Damon Clark and Steve Chang for their comments on this work. We are thankful to the Cognitive and Neural Computational Lab at Yale for their feedback throughout this project, and to Yale Center for Research and Computing for maintaining the HPCs utilized by this project (Misha, Milgram). This work was supported by National Science Foundation (under CAREER Award No. BCS-2441520).

## Supplementary Material

### 1. Details of Programming the Reservoir Computer

This section derives the programming construction of ref. [35] in the notation of the main text, in full generality first, then specialized to the choices used in this paper.

#### 1.1 Objects and Notation

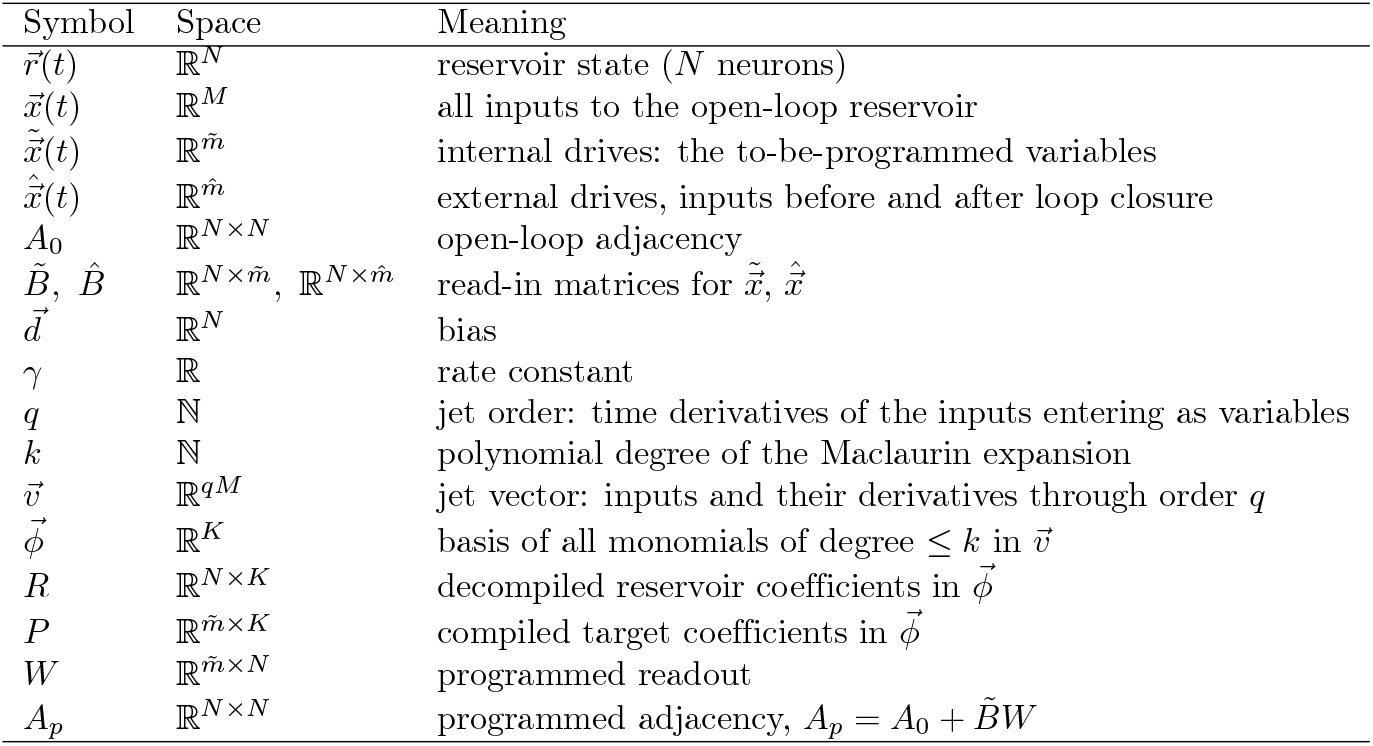

| Symbol | Space | Meaning |
|---|---|---|
| $\vec{r}(t)$ | ℝ<sup>*N*</sup> | reservoir state (*N* neurons) |
| $\vec{x}(t)$ | ℝ<sup>*M*</sup> | all inputs to the open-loop reservoir |
| $\tilde{\vec{x}}(t)$ | $\mathbb{R}^{\tilde{m}}$ | internal drives: the to-be-programmed variables |
| $\hat{\vec{x}}(t)$ | $\mathbb{R}^{\hat{m}}$ | external drives, inputs before and after loop closure |
| *A*<sub>0</sub> | ℝ<sup>*N*×*N*</sup> | open-loop adjacency |
| $\tilde{B},\ \hat{B}$ | $\mathbb{R}^{N\times\tilde{m}},\ \mathbb{R}^{N\times\hat{m}}$ | read-in matrices for $\tilde{\vec{x}}$, $\hat{\vec{x}}$ |
| $\vec{d}$ | ℝ<sup>*N*</sup> | bias |
| *γ* | ℝ | rate constant |
| *q* | ℕ | jet order: time derivatives of the inputs entering as variables |
| *k* | ℕ | polynomial degree of the Maclaurin expansion |
| $\vec{v}$ | ℝ<sup>*qM*</sup> | jet vector: inputs and their derivatives through order *q* |
| $\vec{\phi}$ | ℝ<sup>*K*</sup> | basis of all monomials of degree ≤ *k* in $\vec{v}$ |
| *R* | ℝ<sup>*N*×*K*</sup> | decompiled reservoir coefficients in $\vec{\phi}$ |
| *P* | $\mathbb{R}^{\tilde{m}\times K}$ | compiled target coefficients in $\vec{\phi}$ |
| *W* | $\mathbb{R}^{\tilde{m}\times N}$ | programmed readout |
| *A*<sub>*p*</sub> | ℝ<sup>*N*×*N*</sup> | programmed adjacency, $A_p = A_0 + \tilde{B}W$ |

The inputs decompose as 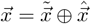 with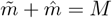. The tilde variables are internal driving forces with a destiny: after programming, the closed-loop reservoir generates them itself through the feedback 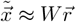. The hat variables remain genuine inputs throughout. **In this paper**, the dynamical algorithm has 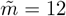 variables and there are no external drives during simulation, 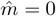 (inputs are used only to set initial conditions; see Initialize and Simulate below), so the programmed reservoir is fully autonomous.

#### 1.2 The Open-Loop Reservoir

The open-loop reservoir is the driven system

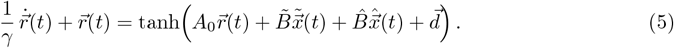

We expand its response about a joint operating point 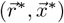 of the state and the inputs. Requiring 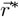 to be a fixed point of Eq. 5 at constant input 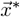 determines the bias:

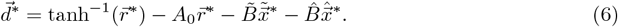

##### Choices used here

*A*_0_ = 0 (the open-loop reservoir has no recurrence of its own; all recurrence in *A*_*p*_ is programmed), the input operating point at the origin, 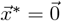, and 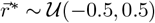 drawn randomly. Under these choices Eq. 6 reduces to 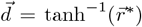, and 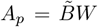. The read-in matrices 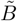, 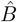 are drawn i.i.d. from *U*(−0.5, 0.5).

#### 1.3 The Jet Vector and the Shared Basis

Decompilation and compilation meet in a single coordinate system: a polynomial basis over the inputs and their time histories. Fix a jet order *q* and collect the inputs and their time derivatives through order *q* into the jet vector

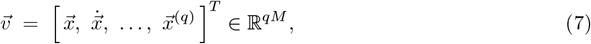

noting that 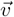 contains the jets of both the internal and the external drives. The shared basis 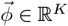 consists of all monomials of degree at most *k* in the components of 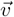, including the constant, so that

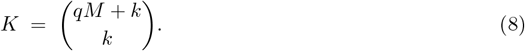

##### Choices used here

*q* = 1 (inputs and their first derivatives) and *k* = 3 (cubic Maclaurin), so with 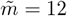 and 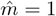 for the paddle input but was kept constant and out of board interactions,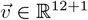 and 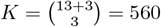.

#### 1.4 Decompile the Reservoir into the Basis

The programming method rests on a symbolic closed-form approximation of the driven reservoir’s response about the operating point. Both sides of Eq. 5 are useful: the right-hand side 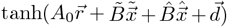 equals the instantaneous update 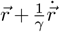 exactly, and the closed-form solution expresses this update as a smooth map *h* of the jet vector,

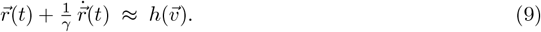

Expanding *h* as a multivariate Maclaurin series of degree *k*,

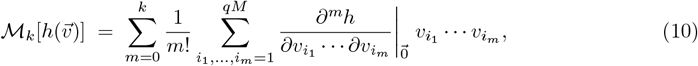

collects the reservoir’s expansion coefficients into the decompiled matrix

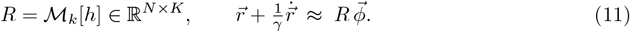

The expansion is a derivational object: it is used once, to obtain *R*, and no expanded or reduced system is ever simulated.

#### 1.5 Compile the Dynamical Algorithm into the Basis

The dynamical algorithm specifies the internal variables’ flow field,

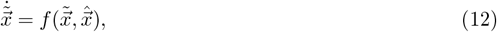

where the component functions *f*_*i*_ may be constant registers (*f*_*i*_ = 0), linear, polynomial (e.g., the cubic bifurcation 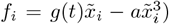), or coupled across variables. The compilation target pairs each side of Eq. 11 with its algorithmic counterpart. The reservoir’s instantaneous update 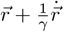 corresponds, through the readout, to the algorithm’s instantaneous update 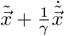, so the target is

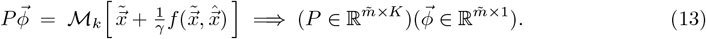

#### 1.6 Program the Readout

Programming is the least-squares match of the two compiled objects in the shared basis:

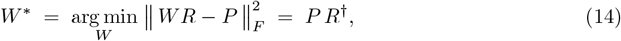

with *R*^*†*^ the Moore–Penrose pseudoinverse. Matching coefficients matches functions, 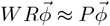, which by Eqs. 11 and 13 reads

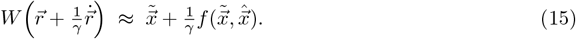

Equation 15 carries both guarantees at once: its zeroth-order content is readout consistency, 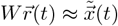, and its derivative content is the dynamics, 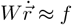. This is the sense in which the reservoir’s latent flow is semiconjugate to the target flow [40, 41].

#### 1.7 Close the Loop

‘Substituting the readout for the internal drives, 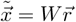, eliminates the programmed inputs and yields the programmed reservoir,

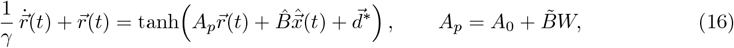

which, under the choices used here 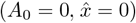, is the autonomous system 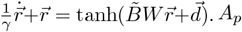 is dense and typically has a spectral radius ≫1; the system is this fully nonlinear network of leaky integrators.

#### 1.8 Initialize and Simulate

To simulate a specific Pong condition:

1. Set the trial’s initial configuration 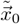 (ball position and velocity, plus the algorithm’s constant registers) through the input channel of the open-loop reservoir and evolve to convergence, 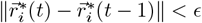, at the conditional operating point: the fixed point

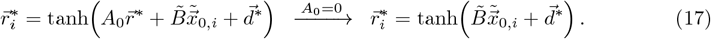

Each conditional operating point indexes a unique board configuration, displaced from the global operating point in the *N*-dimensional state space.
2. Numerically integrate the programmed reservoir from 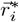 (Runge–Kutta 4).
3. Read out the algorithm’s variables at any time as 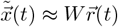.

#### 1.9 Hyperparameters Used in This Paper

| Parameter | Value | Notes |
|---|---|---|
| *N* | 1000 | reservoir units |
| *m̃* | 12 | dynamical-algorithm variables |
| *m̂* | 1 | disconnected paddle drive during simulation |
| *A*<sub>0</sub> | 0 | all recurrence programmed |
| *q* | 1 | jet order |
| *k* | 3 | Maclaurin degree |
| γ | 100 | rate constant |
| ϵ | 10<sup>−13</sup> | convergence tolerance for loading |
| solver | Runge-Kutta 4 | dt = 0.01 |

## 2 The Lotka–Volterra System

The main text applies dSPM to a single dynamical algorithm, the Pong gameboard. Here we program a second, textbook system into a reservoir with the same six-step procedure, both to illustrate our interface for building dSPMs and to test quantitatively how faithfully a programmed reservoir preserves the structure of its target. The Lotka–Volterra model of predator-prey dynamics is governed by

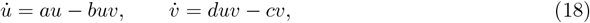

where *u* and *v* denote the prey and predator populations and *a, b, c, d >* 0 are rate constants (written in Latin letters to avoid confusion with the reservoir’s rate constant, *γ*). Its orbits are closed and nested about a single center, the coexistence fixed point (*c/d, a/b*), and each conserves the quantity

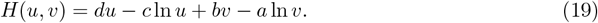

Three properties make this system a demanding test of dSPM: its non-linearity is not separable, since the *uv* coupling terms require multivariate polynomial basis functions of order ≥ 2; its conserved quantity is exact in the true system, so that any deviation along a reservoir trajectory reflects error in the embedded flow field; and, unlike the Pong gameboard, its programming was not the focus of development or tuning.

Supplementary Fig. 1 shows the entire procedure in our Matlab-based interface, in less than 60 lines: the dynamical algorithm is specified symbolically, a reservoir is initialized, and a three-line compile sequence (`decompile, program, close loop`), corresponding to Steps 2 through 6 of the Methods and identical to the one used for the Pong gameboard, programs it. Run from seven initial conditions spanning a wide range of orbital amplitudes, the programmed reservoir reproduces the closed, nested orbits of the Lotka–Volterra system about its coexistence fixed point, with the correct winding direction and without systematic distortion (Supplementary Fig. 2, left and right panels). Over 20,000 time steps, *H* remains within 3 × 10^−5^ of its initial value on every orbit in each of five reservoirs, with a median deviation of 3 ×10^−6^ on the largest orbit (Supplementary Fig. 2, middle panel). This tolerance is not intrinsic to the method. Halving the input scale from 0.02 to 0.01 reduced the deviation more than thirtyfold, and it stops improving only where the readout meets the limits of double precision. The programmed reservoir thus preserves the invariant structure of its target quantitatively, and not only its qualitative topology; a reservoir with random, unprogrammed connectivity has no such property, whatever its spectral radius.

This example shows how cognitive science and neuroscience modelers can express a dynamical algorithm symbolically, initialize a reservoir with a desired number of units, and program it to obtain a topologically semiconjugate recurrent network, without training. It also demonstrates the expressivity of dSPM’s reservoirs beyond the Pong gameboard: a non-linear, conservative system that played no part in the development of the framework is embedded with its invariant structure intact, suggesting that dSPMs can generalize to a broad class of dynamical algorithms that correspond to structure-preserving representations for complex domains of cognition. The system is also of direct neurobiological interest. Its core motif, a self-amplifying variable held in check by a second variable that it in turn sustains, recurs in classical models of neural dynamics: in the excitation and recovery variables of the FitzHugh–Nagumo model of the single neuron [88], and in the interaction of excitatory and inhibitory populations [89]. Rate equations of Lotka–Volterra type also arise directly as descriptions of competing neurons and neuronal populations [90, 91]. How a programmed reservoir achieves this, in terms of its connectivity and of its stability at each point of a trajectory, is the subject of Supplementary Section 3, where we show that the conserved quantity has a direct counterpart in the reservoir’s spectrum.

**Supplementary Figure 1.**
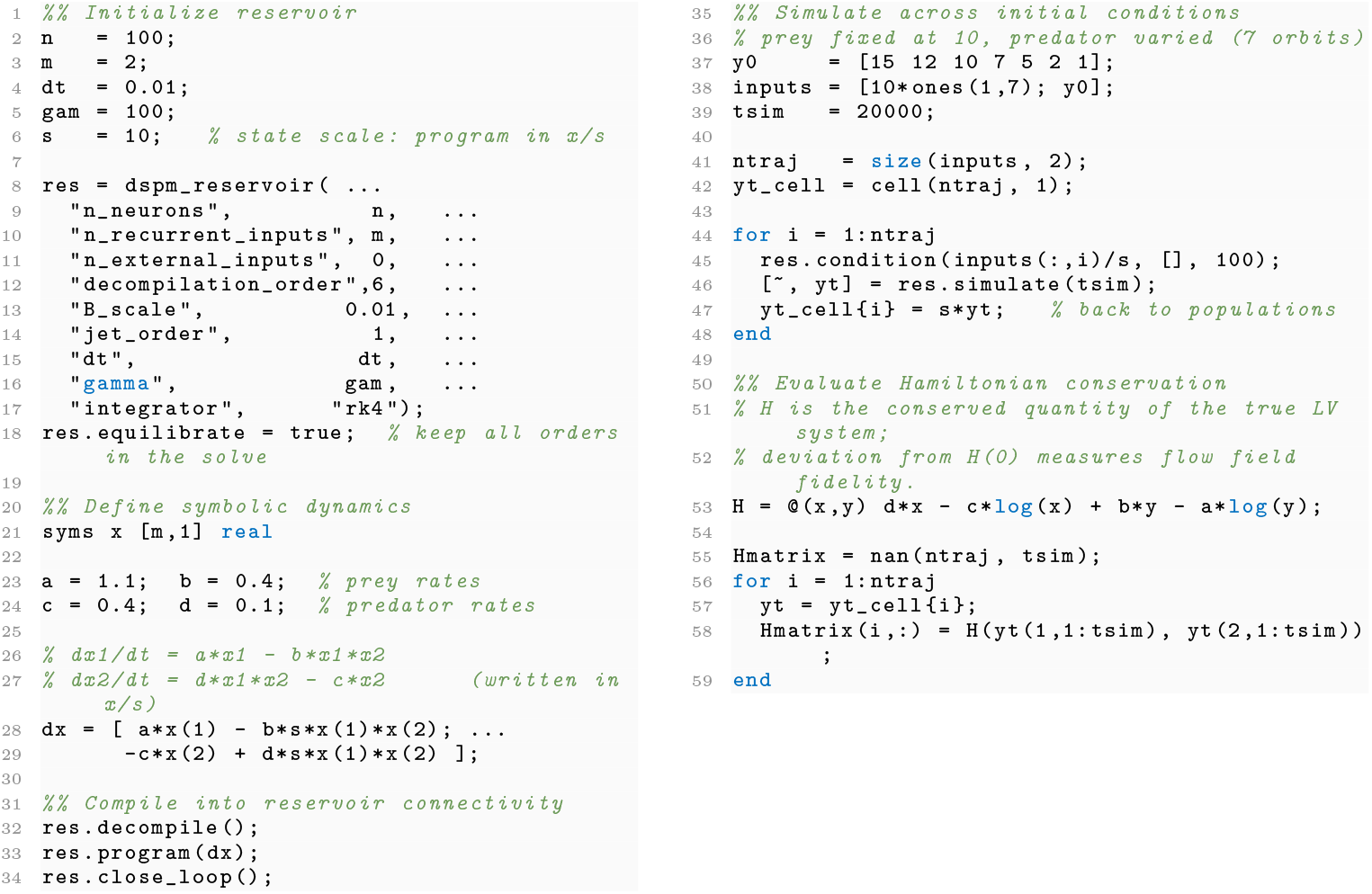
Analytical programming of Lotka–Volterra dynamics into a dSPM reservoir. The three-line compile sequence (`decompile, program, close_loop`) is identical to the procedure used for the Pong gameboard. Seven initial conditions are simulated, with the prey population fixed and the predator population varied, and the deviation of the Hamiltonian from its initial value along each trajectory is used as a quantitative criterion for flow-field fidelity.

**Supplementary Figure 2.**
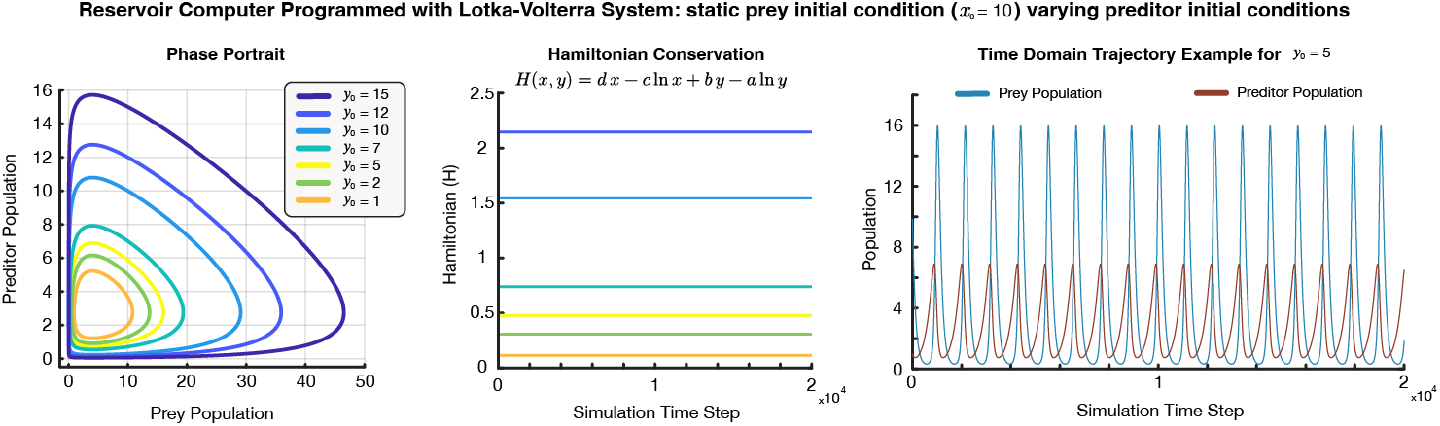
Demonstrating the computational expressivity and power of the dSPM framework. The Lotka–Volterra predator-prey system (*a* = 1.1, *b* = 0.4, *c* = 0.4, *d* = 0.1) was embedded into a reservoir computer of *N* = 100 units using the six-step analytical programming procedure described in the Methods and implemented in Supplementary Fig. 1, with the prey initial condition fixed at *x*_0_ = 10 and the predator initial condition varied across seven values. No parameters were fit to the reservoir’s dynamics. (Left) Phase portrait of the programmed reservoir’s readout *W r*(*t*) for all seven initial conditions. (Middle) Hamiltonian *H*(*x, y*) = *d x* − *c* ln *x* + *b y* − *a* ln *y* evaluated along each reservoir trajectory over 20,000 time steps. (Right) Time-domain trajectory for *y*_0_ = 5, showing stable periodic oscillations of the prey (blue) and predator (red) populations with no drift or amplitude modulation. The spectral properties of this reservoir are reported in Supplementary Section 3.

## 3 Spectral Properties of dSPM Reservoirs

Here we provide the evidence for the statements in the Methods about the spectral radius of the programmed adjacency, using the same 50 Pong reservoirs analyzed in the main text. We report two quantities: the conventional spectral radius, *ρ*(*A*_*p*_) [25, 42], and the spectrum of *A*_*p*_ after accounting for the gradient of tanh at the reservoir’s current state, diag(1 − *r*^2^) *A*_*p*_, which is the Jacobian of the reservoir’s tanh map at its working point [35, 92]. For the latter, a value of 1 corresponds to a state that is ‘held’ (the recurrent input exactly offsets the leak), and a mode grows when the real part of its eigenvalue exceeds 1; we refer to the eigenvalue with the largest real part as the *leading eigenvalue*.

### At its operating points, the spectrum is that of the dynamical algorithm

Across the 50 reservoirs, *ρ*(*A*_*p*_) ranges from 3.4 to 5.0 × 10^3^, varying with each reservoir’s random *B* and *r*^*^ (Supplementary Fig. 3A). Evaluated at the analytic operating point, however, the spectral radius is 1.20 in all 50 reservoirs, and its eigenvalues coincide with those of the first-order term of the target’s expansion, *P*. The largest belongs to the switch that stores the ball’s direction of travel. At each trial’s conditional operating point, 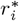 the value is 1.00 for all 79 conditions in all reservoirs: the reservoir holds that trial’s configuration without decay. The large *ρ*(*A*_*p*_) arises from the nonlinear terms of the expansion, which carry the switches and collision detectors: solving for *W* against the first-order terms alone gives a spectral radius of 1.3 in each of 50 reservoirs generated with the same settings. The Lotka–Volterra reservoir of Supplementary Section 2 obeys the same relation. Its eigenvalues are 1 + *a/γ* and 1 − *c/γ* at the origin, reproducing the saddle of the target, and 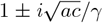 at the coexistence fixed point, where a real part of 1 is the local counterpart of the conserved Hamiltonian; changing the scale of *B* alters *ρ*(*A*_*p*_) and leaves these adjusted eigenvalues unchanged.

### Along a condition, the reservoir leaves its ‘held’ regime only at a collision

Evaluated at every integration step over the duration of each condition, the leading adjusted eigenvalue remains at 1 throughout conditions without a bounce. In conditions with a bounce, it departs from 1 twice (Supplementary Fig. 3B). It first rises linearly as the ball passes the threshold of a collision detector, and returns to 1 once the detector switches on. Then, as the detector drives the direction switch through its midpoint, it rises sharply and briefly, peaking at 1.10 in all 50 reservoirs for approximately 40 ms (Supplementary Fig. 3C) and never exceeding 1.12. The slope of the first rise is proportional to the ball’s speed toward the wall, |*v*_*y*_|, across the 26 bounce conditions (*r* = 0.99; Supplementary Fig. 3D) — during this period, the spectrum of the reservoir tracks a variable of the simulated scene.

### Implications

These results make three points about dSPM’s programmed reservoirs. First, the spectral radius of *A*_*p*_ is a property of a particular random reservoir, whereas the spectrum at its operating points is a property of the dynamical algorithm: fifty different reservoirs share it to numerical precision. Second, a reservoir that is strongly coupled by the conventional measure nonetheless holds its state, and departs from this regime only at the moments the algorithm requires a nonlinear computation, namely detecting a collision and reversing the ball’s direction. A regime that traditional reservoir computing avoids [25, 86] is here a controlled part of the computation. Third, these departures are themselves structured by the scene being simulated, as the dependence on the ball’s speed shows. Together, they illustrate concretely what it means for the reservoir to be topologically semiconjugate to its target: the reservoir’s own stability properties, at each moment, are those of the dynamical algorithm. They also show what the programming procedure makes possible. It does not avoid instability; it uses instability. A regime of strong coupling and marginal stability that conventional reservoir design excludes is here specified analytically, without training, and each departure from stability carries out a step of the computation.

**Supplementary Figure 3.**
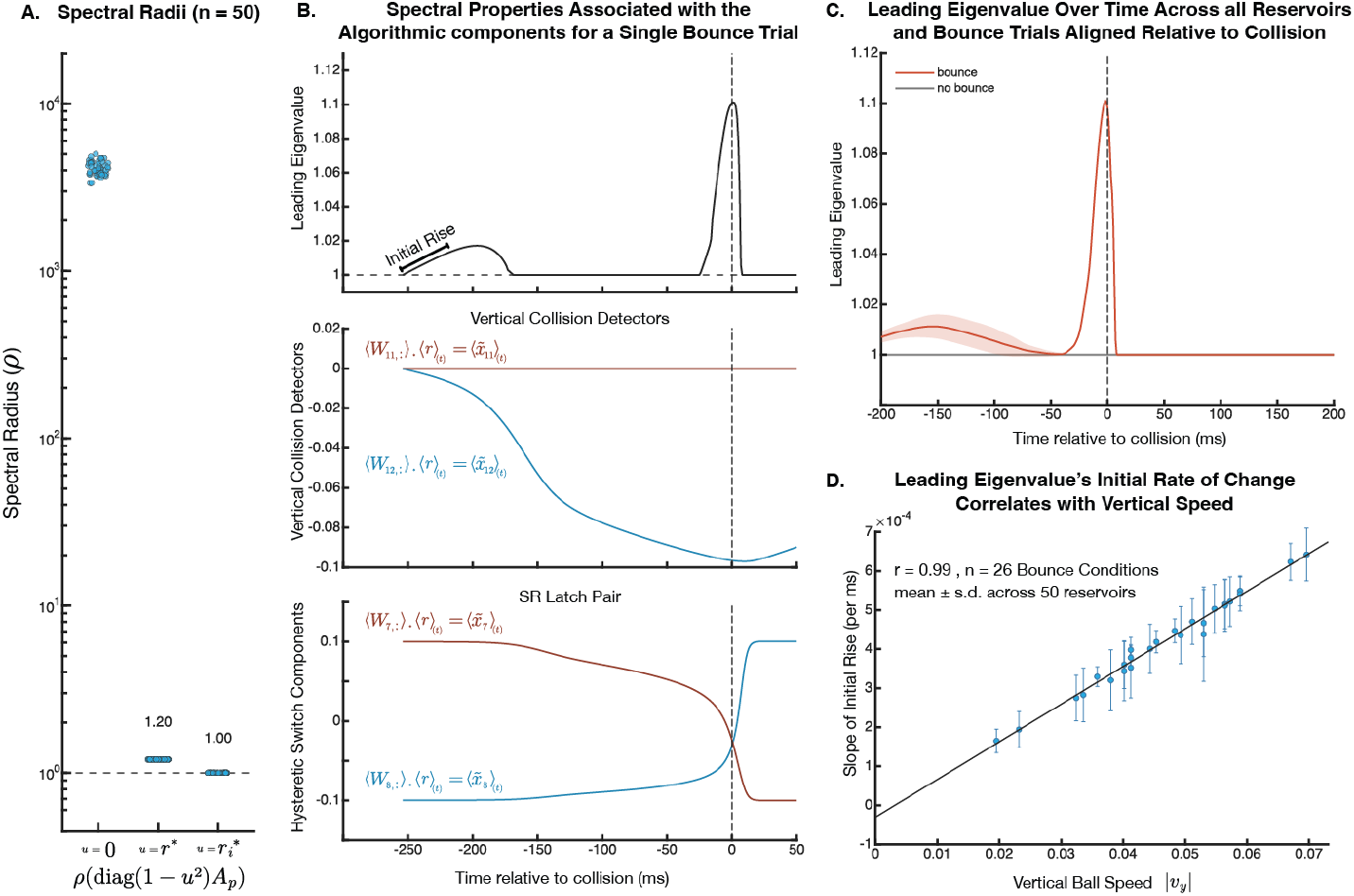
Spectral properties of dSPM’s programmed reservoirs. (A) Three readings of the same connectivity for each of the 50 Pong reservoirs (*N* = 1000): the spectral radius of the programmed adjacency, *ρ*(*A*_*p*_); the spectral radius after accounting for the gradient of tanh at the analytic operating point *r*^*^; and the same at the conditional operating points 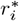, the state of the reservoir at *t* = 0 once it has been conditioned on the initial configuration of condition *i* (mean over the 79 conditions). Each dot is one reservoir; note the logarithmic axis. *ρ*(*A*_*p*_) varies across reservoirs, whereas the value at *r*^*^ is the same in all of them. Dashed line, 1. (B) A one-bounce condition from one reservoir, aligned to the bounce (dashed line). Top, leading adjusted eigenvalue (largest real part) evaluated at the reservoir’s state at each integration step. Middle, the two collision detectors for the top and bottom walls. Bottom, the pair of switch variables that store the ball’s vertical direction. Each of these traces is a single row of the programmed readout *W* applied to the full state of the densely connected reservoir, 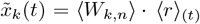; no unit is dedicated to any variable. The leading eigenvalue first rises as the ball passes a detector’s threshold, returns to 1 when the detector switches on, and rises again briefly as the switch flips. (C) Leading eigenvalue aligned to the bounce, averaged over bounce conditions (red) and, at matched times, over no-bounce conditions (gray). Lines and shading, mean ± s.d. across the 50 reservoirs. (D) Slope of the initial rise in (B) against the ball’s speed toward the wall (the magnitude of its vertical velocity, |*v*_*y*_|, read out through *W* before the bounce), for each of the 26 one-bounce conditions. Points and error bars, mean *±* s.d. across the 50 reservoirs. Line, least-squares fit.

## 4 Pseudo-Code of the Board Simulation

**Supplementary Figure 4.**
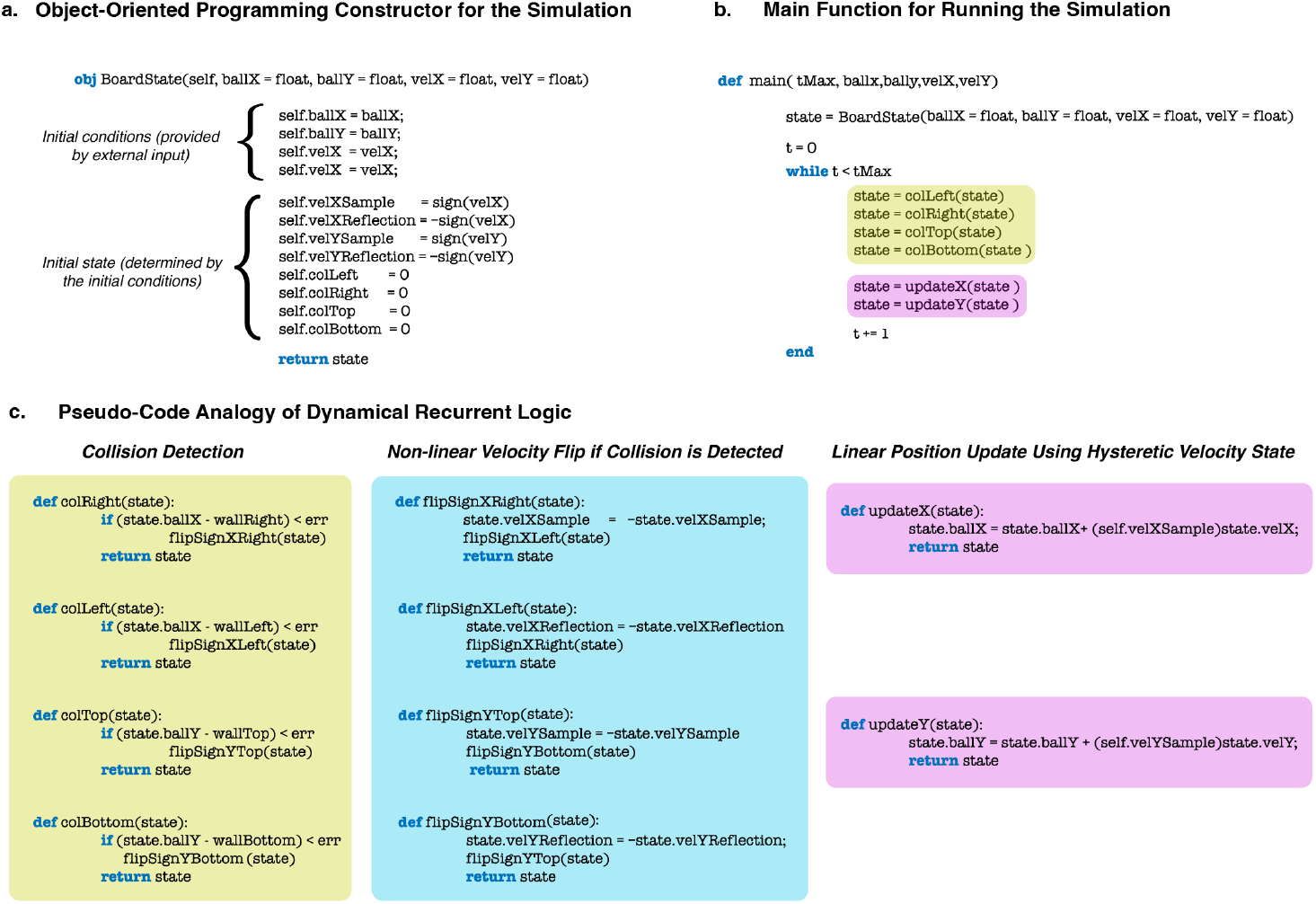
Pseudocode of the physics-based representation of the gameboard, which underlies our dSPM construction. This figure presents an intuitive pseudocode representation to illustrate how the 12 coupled differential equations (Equations 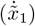 through 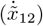) described in the Methods. (A) An object-oriented programming constructor of the board state, showing how initial conditions map to the initial (*x, y*) position of the ball within the game board and the constant velocity registers (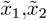 from equations 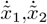 in Methods) and initialize the hysteretic state variables. (B) Main loop for unfolding the board dynamics. (C) Pseudocode modules corresponding to the computational blocks 2-4 of the Methods section: Collision Detection (yellow) represents the dynamics of 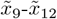 (Equations 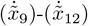 in Methods), which implement threshold-based collision detection; Non-linear Velocity Flip (blue) illustrates the hysteretic NAND gate dynamics of 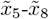 (Equations 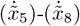 in Methods) that maintain velocity sign memory through bistable attractors; Linear Position Update (purple) shows how 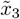 and 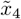 (Equations 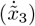 and 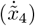 in Mehtods) integrate velocity to update position. These functions make recurrent calls to each other. In dSPM, this physics-based representation is analytically embedded into the connectivity matrix of a reservoir computer, 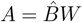.

## 5 Dynamical algorithm as a low dimensional set of symbolic ODEs

**Supplementary Figure 5.**
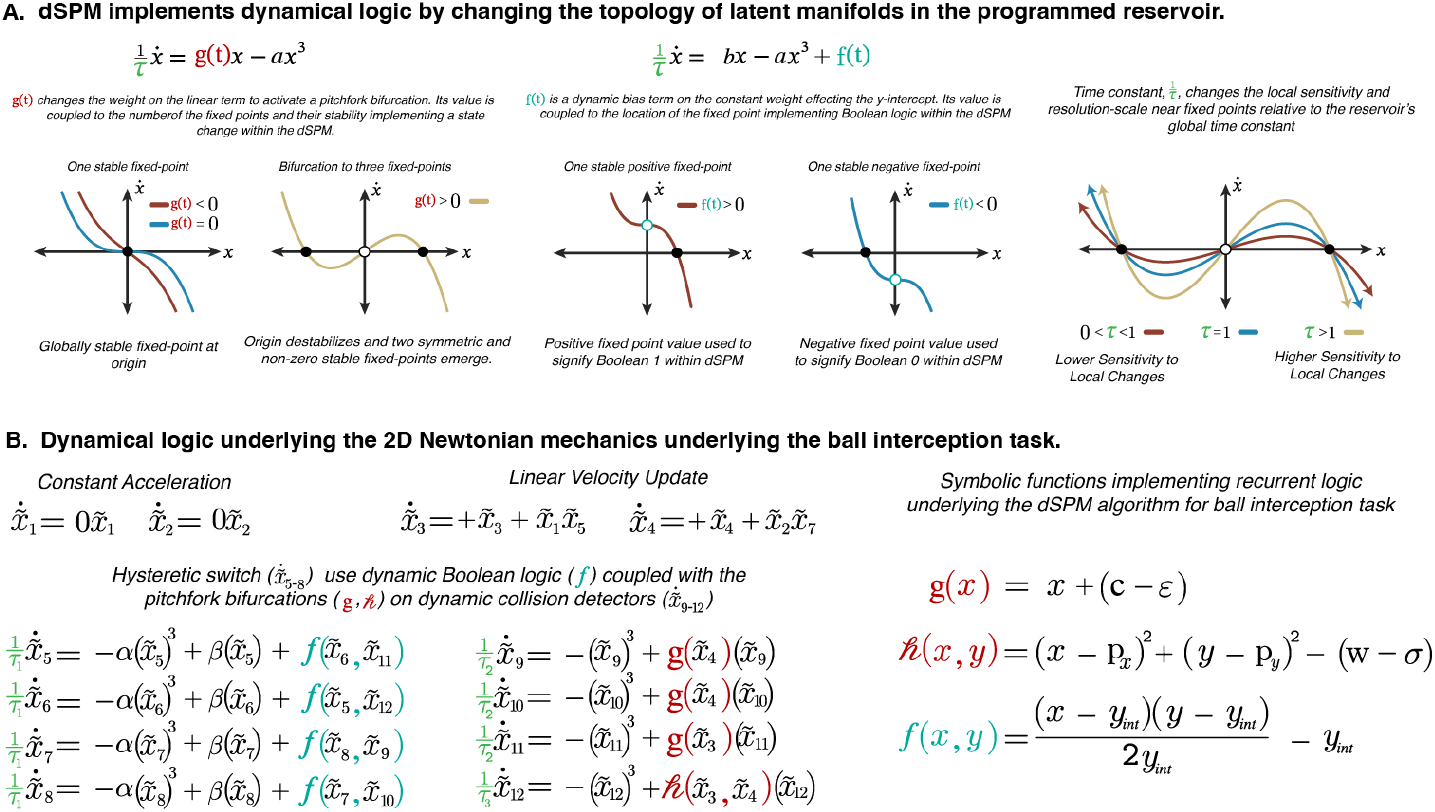
Example dynamical primitives for implementing symbolic representations. (A) Left two panels show the phase portraits of a controlled bifurcation with the equation 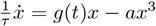. The stable fixed point (attractor; filled circle) on the left transitions into two stable fixed points and one unstable fixed points (repeller; unfilled circle) as *g*(*t*) becomes positive. We use this type of bifurcation for collision detection in our gameboard representation. Middle two panels show the same type of bifurcation but this time with its constant term time-varying, 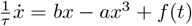. The stable and unstable fixed points non-linearly transition as a function of the time-varying constant term *f*(*t*). In our gameboard representation, we use this type of bifurcation for the velocity sign variable. Through these time-varying coefficients (e.g., *g*(*t*) and *f*(*t*)), we can couple different dynamical variables and implement branching logic, linear velocity dynamics, and other elements needed to represent the physical scenes and other structure-preserving representations. The rightmost panel shows how the sensitivity of these dynamical primitives can be changed by adjusting the value of *τ* . This rightmost panel plots the second plot from the left (i.e., 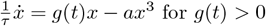) with different values of the time constant. (B) The full dynamical algorithm, consisting of the 12 dynamical variables, as described in Methods.

## 6 Animation of dSPM’s Dynamical Algorithm and Reservoir Computer State

**Supplementary Figure 6.**
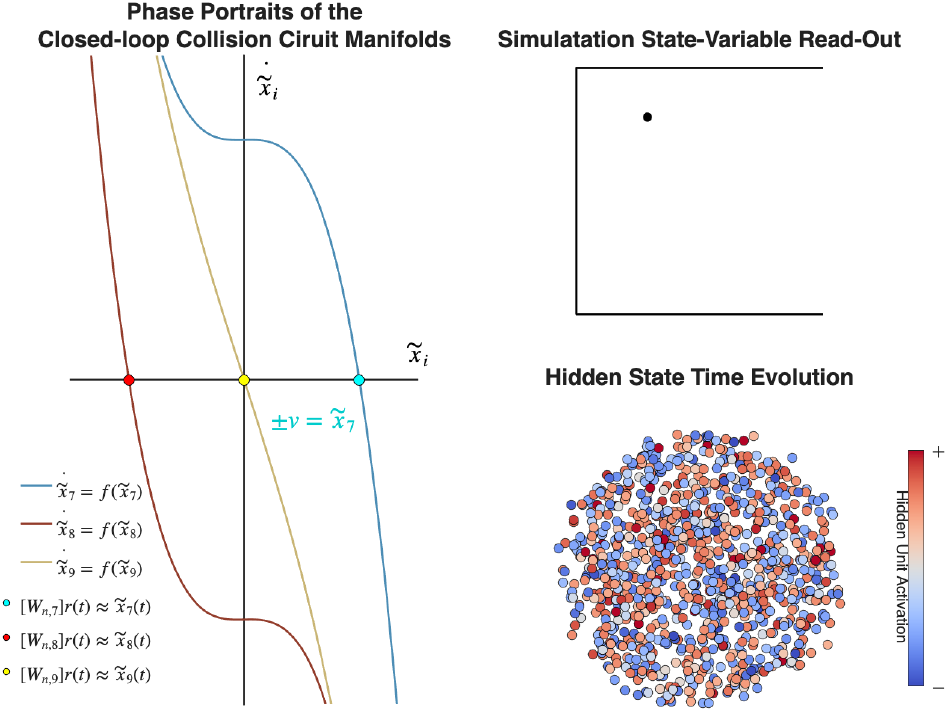
Animated visualization of the dSPM dynamical algorithm and reservoir state evolution (1000 units) implementing non-linear wall collision detection and resolution. Left panel: Phase portraits of the collision circuit’s recurrent dynamics, showing the theoretical nullclines (continuous curves) programmed via the polynomial-shift operators and pitchfork bifurcations described in Methods (equations 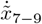, the two SR latch components and the top wall collision detector respectively) which recurrently couple to 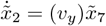, the symbolic representation of the ball’s vertical velocity which integrates to its position 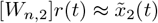. The points represent the actual instantaneous values of these dynamical variables as read out from the network state via 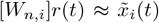. The blue and red curves show the hysteretic switch’s pair-dynamics (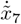 and 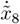 for vertical velocity), implementing bistable memory. The yellow curve represents the collision detector, which transitions between stable zero and non-zero fixed points when the ball approaches the boundary. The cross-coupling between collision detection and velocity sign variables implements the NAND gate logic triggering velocity reversal upon collision. Top right panel: Real-time ball trajectory on the board, computed from the linear position update equations 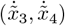 that integrate the velocity components stored in the hysteretic switches. The red trace shows the ball’s path. The wall collision triggers a transition in the dynamical landscape (left panel), flipping the appropriate velocity component while maintaining the orthogonal component unchanged. Bottom right panel: High-dimensional hidden state of PAN showing all *N* = 1000 neuron activations *r*(*t*) ∈ R^*N*^ at each time point. The color gradient (from dark blue to light cyan) encodes individual neuron activities. Despite this high-dimensional embedding, the network autonomously constrains its dynamics to the 12-dimensional manifold specified by the programmed differential equations. The *L*_2_-norm solution that creates the programmed adjacency 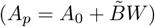 during the compilation step, distributes the low-rank symbolic/dynamical representations heterogeneously across the population-rank of the network. The relatively uniform distribution of activities indicates that the network utilizes its full representational capacity rather than sparse coding, consistent with such a *L*_2_-norm based analytical encoding.

NOTE: To play the animation within the PDF, please use Adobe Acrobat. This animation is also included as a Supplementary Video.

## 7 Accounting for DMFC’s noise floor in the dimensionality analyses

DMFC responses are averages over a finite number of trials and so inherently involve a noise floor that the deterministic models do not. Noise spreads variance across many dimensions, so it raises DMFC’s participation ratio. We examined how much of the difference between DMFC and dSPM in Fig. 2B this accounts for by adding DMFC-matched noise to dSPM.

**Supplementary Figure 7.**
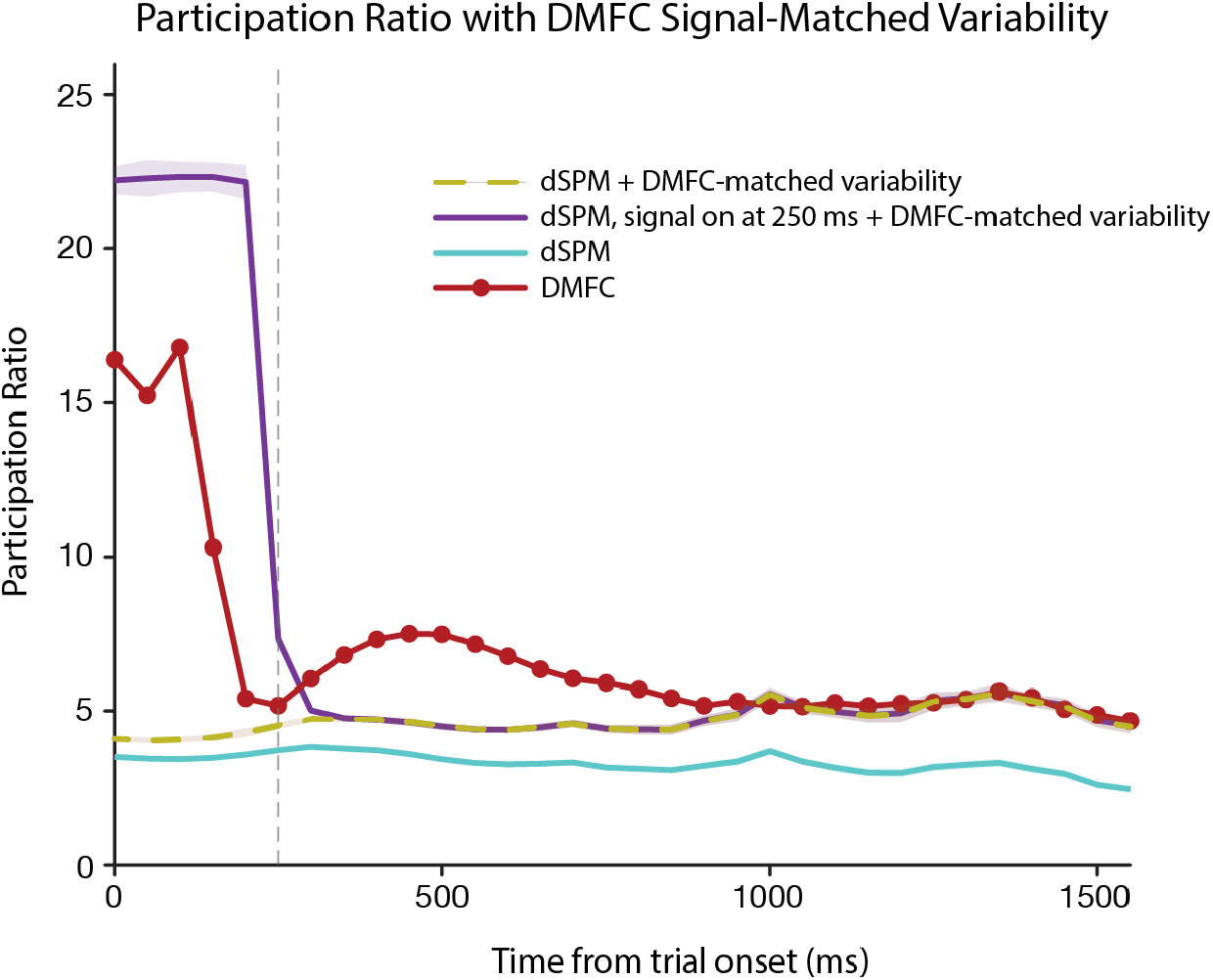
The gap between dSPM and DMFC’s time-resolved effective dimensionality largely closes after accounting for their different variability profiles. In the yellow curve, we inject additive i.i.d. Gaussian noise into the dSPM reservoir units with a standard deviation that equates the variability profile of the dSPM units to that of the DMFC neurons. This additive noise uniformly elevates the PR as expected, largely closing the gap post-250 ms, with an especially close match later in the trials. In the purple curve, we gate the contribution of the dSPM units such that early on, it is purely the Gaussian noise (so zero signal), and then, sigmoidally, at 250 ms, it transitions to full signal (plus the same Gaussian noise). The PR under this gated dSPM time series qualitatively reproduces the neural data.

## 8 Concordance between DMFC, dSPM, and alternative methods during full trajectory decoding at 250 ms post trial onset

**Supplementary Figure 8.**
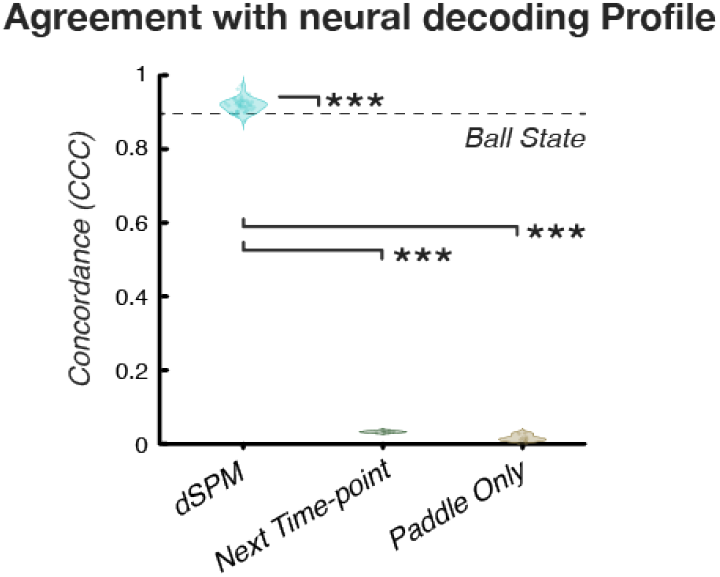
Agreement between model and neural decoding profiles for future-trajectory decoding. For each model instance, the time course of future trajectory decoding accuracy (Fig. 3D) was compared with the corresponding neural (DMFC) time course using Lin’s concordance correlation coefficient (CCC). Unlike Pearson’s r, CCC penalizes offsets and scale differences between the two curves, so a value near 1 indicates that a model reproduces both the shape and the level of the neural profile. Violins show the distribution across instances (dSPM, n = 50 reservoirs; Next Time-point and Paddle-Only, n = 8 trained networks each); points are individual instances. The dashed line marks the single deterministic Ball State readout. dSPM reservoirs matched the neural profile closely (median CCC ≈ 0.92), slightly above the Ball State readout (0.90), while both trained controls were near zero ( ≈0.03 and ≈0.01), reflecting decoding profiles that fall away from the neural curve at longer horizons. Brackets: dSPM vs. each trained control, two-sided exact Wilcoxon rank-sum test; dSPM vs. Ball State, one-sample t-test against the Ball State value as a fixed reference. All p-values are Holm-adjusted across the three comparisons; *** *p <* 0.001.

## 9 Full set of RSA residualization pairs

**Supplementary Figure 9.**
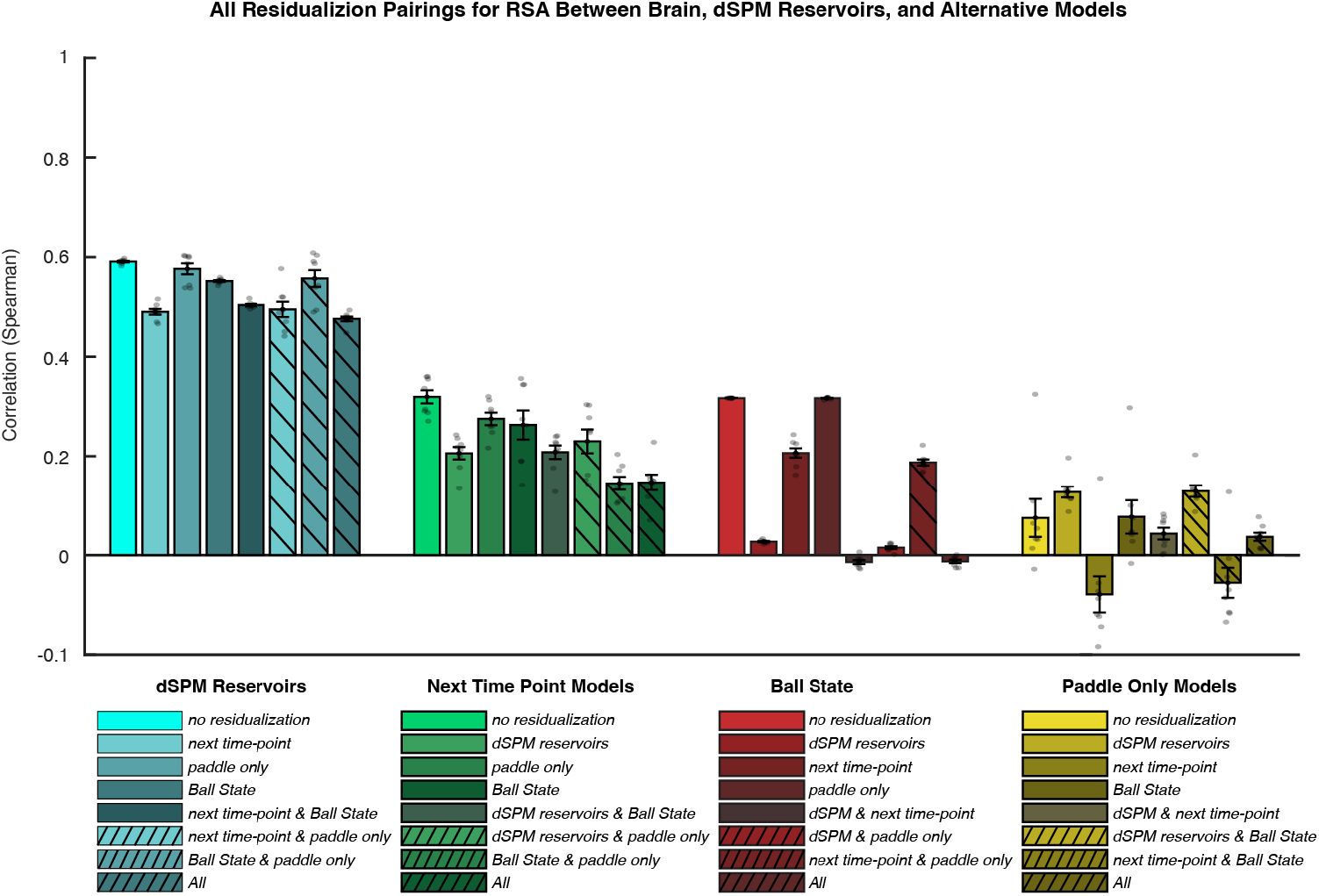
The full set of RSA residualization results. The dSPM model explains more variance than other models and most of this variance is orthogonal to what other models can explain. The reverse is not true: dSPM, for the most part, subsumes what can be explained by the task-optimized RNNs and the task-performant heuristic based on ball state. (For the paddle-only model, because it’s essentially uncorrelated with neural data, our partial regression analysis “pumps in information”, leading to the counterintuitive pattern of increased variance after residualization.)

